# Physiological Adaptability and Parametric Versatility in a Simple Genetic Circuit

**DOI:** 10.1101/2019.12.19.878462

**Authors:** Griffin Chure, Zofii A. Kaczmarek, Rob Phillips

## Abstract

The intimate relationship between the environment and cellular growth rate has remained a major topic of inquiry in bacterial physiology for over a century. Now, as it becomes possible to understand how the growth rate dictates the wholesale reorganization of the intracellular molecular composition, we can interrogate the biophysical principles underlying this adaptive response. Regulation of gene expression drives this adaptation, with changes in growth rate tied to the activation or repression of genes covering enormous swaths of the genome. Here, we dissect how physiological perturbations alter the expression of a circuit which has been extensively characterized in a single physiological state. Given a complete thermodynamic model, we map changes in physiology directly to the biophysical parameters which define the expression. Controlling the growth rate via modulating the available carbon source or growth temperature, we measure the level of gene expression from a LacI-regulated promoter where the LacI copy number is directly measured in each condition, permitting parameter-free prediction of the expression level. The transcriptional output of this circuit is remarkably robust, with expression of the repressor being largely insensitive to the growth rate. The predicted gene expression quantitatively captures the observations under different carbon conditions, indicating that the biophysical parameters are indifferent to the physiology. Interestingly, temperature controls the expression level in ways that are inconsistent with the prediction, revealing temperature-dependent effects that challenge current models. This work exposes the strengths and weaknesses of thermodynamic models in fluctuating environments, posing novel challenges and utility in studying physiological adaptation.

**Significance:** Cells adapt to changing environmental conditions by repressing or activating gene expression from enormous fractions of their genome, drastically changing the molecular composition of the cell. This requires the concerted adaptation of transcription factors to the environmental signals, leading to binding or releasing of their cognate sequences. Here, we dissect a well characterized genetic circuit in a number of physiological states, make predictions of the response, and measure how the copy number of a regulator and its gene target are affected. We find the parameters defining the regulators behavior are remarkably robust to changes in the nutrient availability, but are susceptible to temperature changes. We quantitatively explore these two effects and discuss how they challenge current models of transcriptional regulation.

## Introduction

Cellular physiology is inextricably tied to the extracellular environment. Fluctuations in nutrient availability and variations in temperature, for example, can drastically modulate the cell’s growth rate, which is often used as a measure of the evolutionary fitness (1). In response to such environmental insults, cells have evolved myriad clever mechanisms by which they can adapt to their changing surroundings, many of which involve restructuring their proteome such that critical processes (i.e. protein translation) are allocated the necessary resources. Recent work exploring this level of adaptation using mass spectrometry, ribosomal profiling, and RNA sequencing have revealed that various classes of genes (termed “sectors”) are tuned such that the protein mass fraction of the translational machinery is prioritized over the metabolic and catabolic machinery in nutrient replete environments (2–6). This drastic reorganization is mediated by the regulation of gene expression, relying on the concerted action of myriad transcription factors. Notably, each gene in isolation is regulated by only one or a few components (7). The most common regulatory architecture in *Escherichia coli* is the simple repression motif in which a transcriptional repressor binds to a single site in the promoter region, occluding binding of an RNA polymerase (7, 8). The simple activation architecture, in which the simultaneous binding of an activator and an RNA polymerase amplifies gene expression, is another common mode of regulation. Combinatorial regulation such as dual repression, dual activation, or combined activation and repression can also be found throughout the genome, albeit with lower frequency (9). The ubiquity of the simple repression and simple activation motifs illustrate that, for many genes, the complex systems-level response to a physiological perturbation boils down the binding and unbinding of a single regulator to its cognate binding sites.

Despite our knowledge of these modes of regulation, there remains a large disconnect between concrete, physical models of their behavior and experimental validation. The simple repression motif is perhaps the most thoroughly explored theoretically and experimentally (9) where equilibrium thermodynamic (10–15) and kinetic (16–19) models have been shown to accurately predict the level of gene expression in a variety of contexts. While these experiments involved variations of repressor copy number, operator sequence, concentration of an external inducer, and amino acid substitutions, none have explored how the physiological state of the cell as governed by external factors influences gene expression. This is arguably one of the most critical variables one can experimentally tune to understand the role of these regulatory architectures play in cellular physiology writ large.

In this work, we interrogate the adaptability of a simple genetic circuit to various physiological stressors, namely carbon source quality and growth temperature. Following the aforementioned thermodynamic models, we build upon this theory-experiment dialogue by using environmental conditions as an experimentally tunable variable and determine their influence on various biophysical parameters. Specifically, we use physiological stressors to tune the growth rate. One mechanism by which we modulate the growth rate is by exchanging glucose in the growth medium for the poorer carbon sources glycerol and acetate, which decrease the growth rate by a factor of ≈ 1.5 and ≈ 4 compared to glucose, respectively. We hypothesize that different carbon sources should, if anything, only modulate the repressor copy number seeing as the relationship between growth rate and total protein content has been rigorously quantified (1, 5, 6, 20). Using single-cell time-lapse fluorescence microscopy, we directly measure the copy number of the repressor in each condition. Under a simple hypothesis, all other parameters should be unperturbed, and we can thus rely on previously determined values to make parameter-free predictions of the fold-change in gene expression.

Despite the decrease in growth rate, both the fold-change in gene expression and the repressor copy number remains largely unaffected. We confirm this is the case by examining how the effective free energy of the system changes between carbon sources, a method we have used previously to elucidate parametric changes due to mutations within a transcription factor (15). This illustrates that the energetic parameters defining the fraction of active repressors and their affinity for the DNA are ignorant of the carbon-dependent physiological states of the cell. Thus, in this context, the values of the biophysical parameters determined in one condition can be used to draw predictions in others.

We then examine how variations in temperature influence the transcriptional output. Unlike in the case of carbon source variation, temperature dependence is explicit in our model: the repressor-DNA binding energy and the energetic difference between the active and inactive states of the repressor are scaled to the thermal energy of the system at 37°C. This is defined via the Boltzmann distribution which states that the probability of a state *p*_*state*_ is related to the energy of that state *ε*_*state*_ as

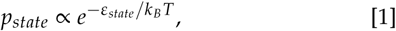

where *k*_*B*_ is the Boltzmann constant and *T* is the temperature of the system. Given knowledge of *T* for a particular experiment, we can easily draw predictions of the fold-change in gene expression. However, we find the fold-change in gene expression is inconsistent with this simple model, revealing an incomplete description of the energetics. We then examine how entropic effects neglected in the initial estimation of the energetic parameters may play an important role; a hypothesis that is supported when we examine the change in the effective free energy.

The results presented here are, to our knowledge, the first attempts to systematically characterize the growth-dependent effects on biophysical parameters in thermodynamic models of transcription. While some parameters of our model are affected by changing the growth rate, they change in ways that are expected or fall close within our *a priori* predictions, suggesting that such modeling can still be powerful in understanding how adaptive processes influence physiology at the level of molecular interactions.

## RESULTS

### Thermodynamic model

We consider a genetic circuit where the expression of a gene is regulated through the binding of an allosteric repressor to the promoter, occluding the binding of RNA polymerase. A thermodynamic rendering of this model, derived previously (13) and in the SI text, computes the fold-change in gene expression relative to an unregulated promoter and has the succinct form

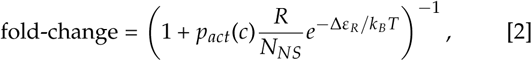

where *R* is the total number of allosteric repressors per cell, *N*_*NS*_ is the number of nonspecific binding sites for the repressor, Δ*ε*_*R*_ is the repressor-DNA binding energy, and *k*_*B*_*T* is the thermal energy of the system. The prefactor *p*_*act*_(*c*) defines the probability of the repressor being in the active state at a given concentration of inducer *c*. In the absence of inducer, *p*_*act*_(*c* = 0) can be written as

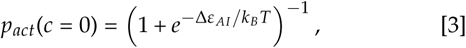

where Δ*ε* _*AI*_ is the energy difference between the active and inactive states. Conditioned on only a handful of experimentally accessible parameters, this model has been verified using the well-characterized LacI repressor of *Escherichia coli* where parameters such as the repressor copy number and DNA binding affinity (10), copy number of the regulated promoter (12, 21), and the concentration of an extracellular inducer (13) can be tuned over orders of magnitude. It has also been shown that this model permits the mapping of mutations within the repressor protein directly to biophysical parameters in a manner that permits accurate prediction of double mutant phenotypes (15). All of these applications, however, have been performed in a single physiological state where cells are grown in a glucose-supplemented minimal medium held at 37°C with aeration. In this work, we challenge this model by changing the environmental conditions away from this gold-standard condition, perturbing the physiological state of the cell.

### Experimental Setup

Seminal studies from the burgeoning years of bacterial physiology have demonstrated a strong dependence of the total cellular protein content on the growth rate (1, 22, 23), a relationship which has been rigorously quantified in recent years using mass spectrometry (2, 3, 5) and ribosomal profiling (6). Their combined results illustrate that modulation of the growth rate, either through controlling the available carbon source or the temperature of the growth medium, significantly alters the physiological state of the cell, triggering the reallocation of resources to prioritize expression of ribosome-associated genes. Eq. 2 has no explicit dependence on the available carbon source but does depend on the temperature through the energetic parameters Δ*ε*_*R*_ and Δ*ε* _*AI*_ which are defined relative to the thermal energy, *k*_*B*_*T*. With this parametric knowledge, we are able to draw quantitative predictions of the fold-change in gene expression in these physiologically distinct states. (3).

We modulated growth of *Escherichia coli* by varying either the quality of the available carbon source (differing ATP yield per C atom) or the temperature of the growth medium [Fig. 1(A)]. All experiments were performed in a defined M9 minimal medium supplemented with one of three carbon sources – glucose, glycerol, or acetate – at concentrations such that the total number of carbon atoms available to the culture remained the same. These carbon sources have been shown to drastically alter growth rate and gene expression profiles (24), indicating changes in the proteomic composition and distinct physiological states. These carbon sources yield an approximate four-fold modulation of the growth rate with doubling times ranging from ≈ 220 minutes to ≈ 65 minutes in an acetate or glucose supplemented medium, respectively [Fig. 1(B) and (C)]. While the growth temperature was varied over 10°C, both 32°and 42°C result in approximately the same doubling time of ≈ 90 min, which is 1.5 times slower than the optimal temperature of 37°C [Fig. 1(B) and (C)].

**Fig. 1.**
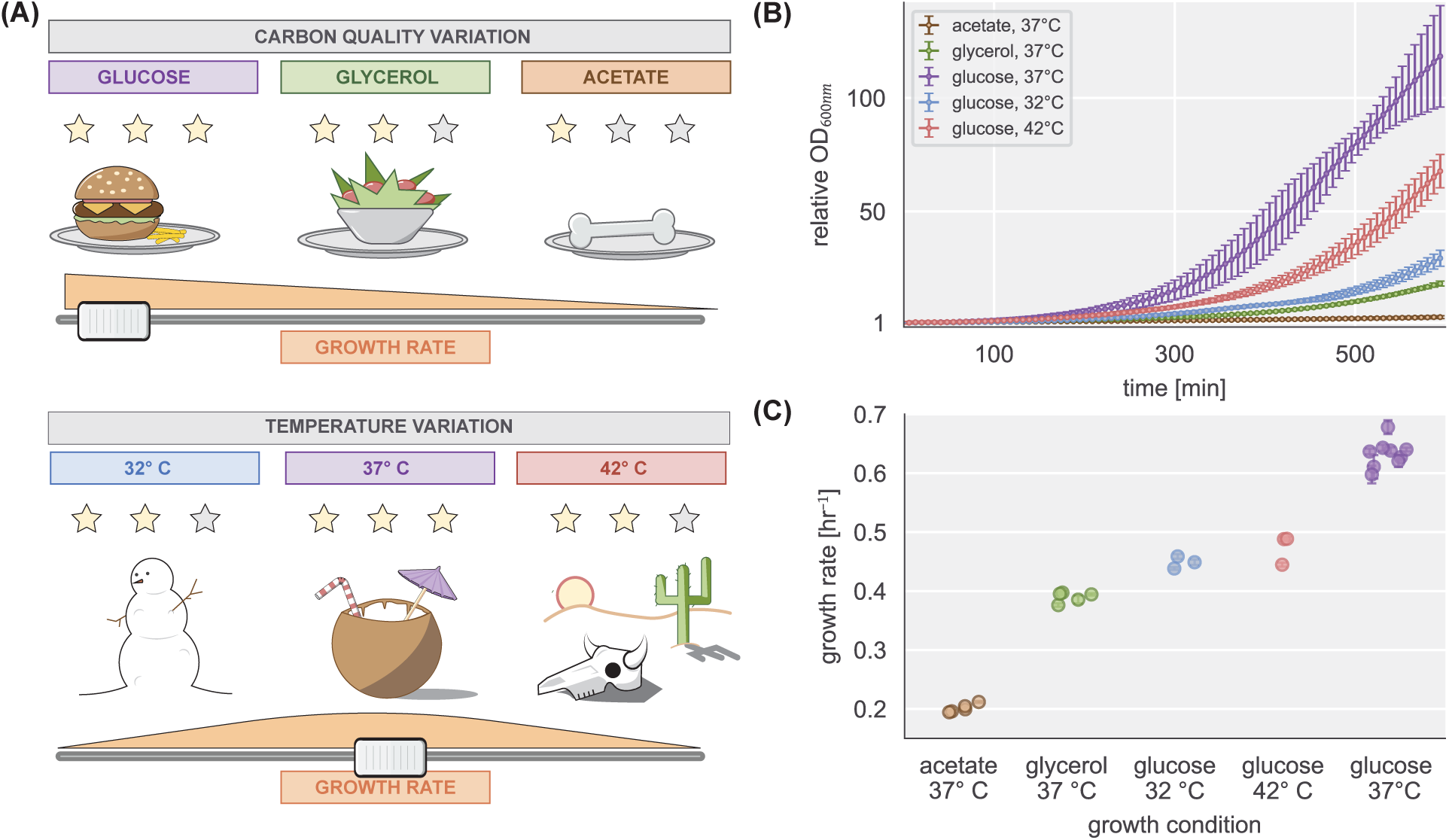
Control of physiological state via growth rate through environmental factors. (A) Bacterial growth can be controlled by varying the available carbon source (top panel) or temperature (bottom panel). (B) Bulk bacterial growth curves under all conditions illustrated in (A). The *y*-axis is the optical density measurements at 600 nm relative to the initial value. Interval between points is ≈ 6 min. Points and errors represent the mean and standard deviation of three to eight biological replicates. (C) Inferred maximum growth rate of each biological replicate for each condition. Points represent the doubling time computed from the maximum growth rate. Error bars correspond to the standard deviation of the inferred growth rate. Where not visible, error bars are smaller than the marker.

The growth rate dependence of the proteome composition suggests that changing physiological conditions could change the total repressor copy number of the cell. As shown in our previous work (13), it can be difficult to differentiate between a change in repressor copy number *R* and the allosteric energy difference Δ*ε* _*AI*_ as there are many combinations of parameter values that yield the same fold-change. To combat this degeneracy, we used a genetically engineered strain of *E. coli* in which the expression of the repressor copy number and its regulated gene product (YFP) can be simultaneously measured. This strain, used previously to interrogate the transcription factor titration effect (12), is diagrammed in Fig. 2(A). A dimeric form of the LacI repressor N-terminally tagged with an mCherry fluorophore is itself regulated through the action of the TetR repressor whose level of activity can be modulated through the addition of the allosteric effector anhydrous tetracycline (ATC). This dual repression genetic circuit allows for the expression of the LacI repressor to be tuned over several orders of magnitude. This is demonstrated in Fig. 2(B) where a titration of ATC in the growth medium results in a steady increase in the expression of the LacI-mCherry gene product (red lines and points) which in turn represses expression of the YFP reporter (yellow lines and points).

**Fig. 2.**
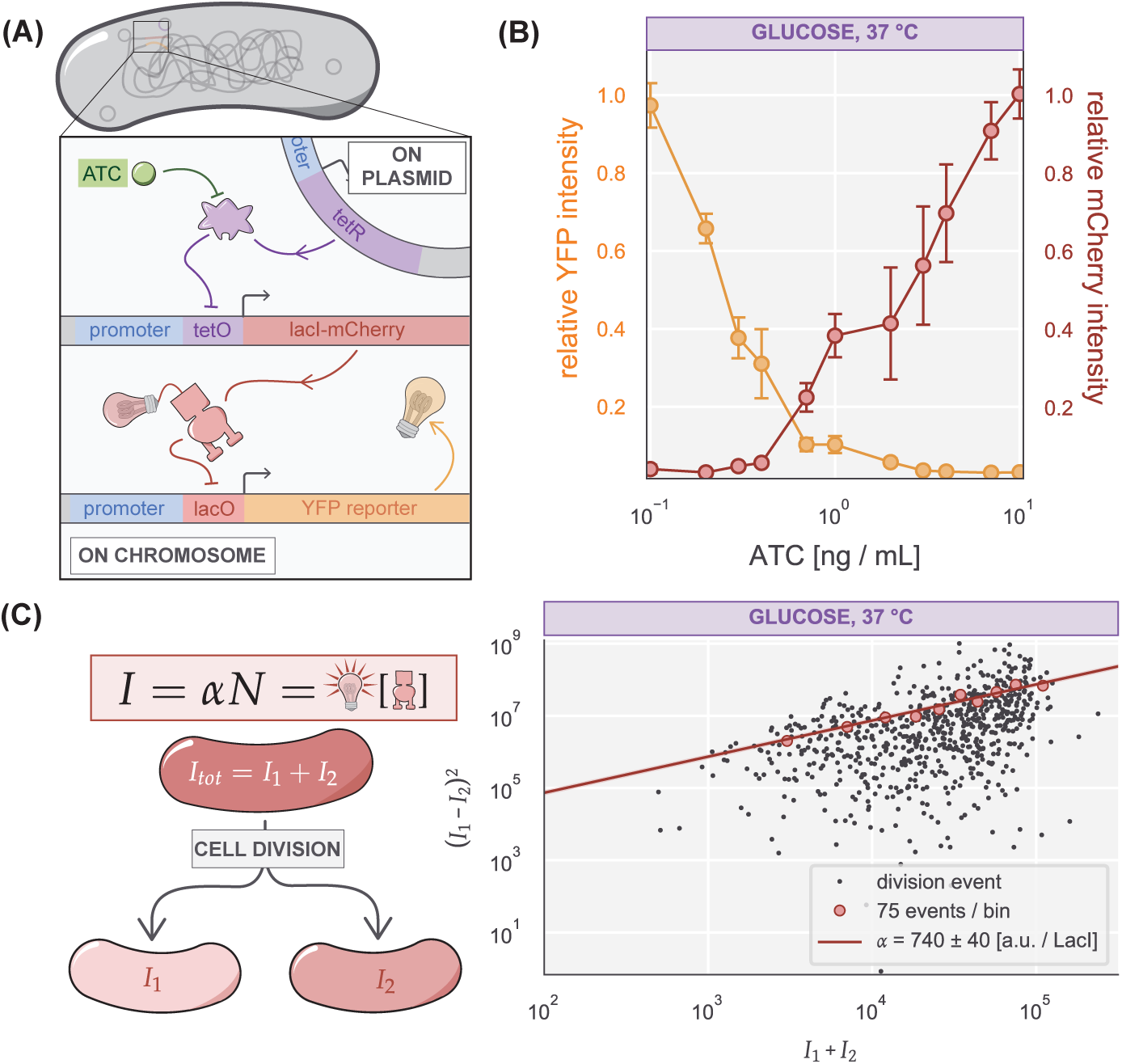
Control and quantification of repressor copy number. (A) The dual repression expression system. The inducible repressor TetR (purple blob) is expressed from a low-copy-number plasmid in the cell and represses expression of the LacI-mCherry repressor by binding to its cognate operator (*tetO*). In the presence of anhydrous tetracycline (ATC, green sphere), the inactive state of TetR becomes energetically favorable, permitting expression of the LacI-mCherry construct (red). This in turn binds to the *lacO* operator sequence repressing the expression of the reporter Yellow Fluorescent Protein (YFP, yellow lightbulb). (B) An ATC titration curve showing anticorrelated YFP (yellow) and mCherry (red) intensities. Reported values are scaled to the maximum mean fluorescence for each channel. Points and errors correspond to the mean and standard error of eight biological replicates. (C) Determination of a fluorescence calibration factor. After cessation of LacI-mCherry expression, cells are allowed to divide, partitioning the fluorescently tagged LacI repressors into the two daughter cells (left panel). The total intensity of the parent cell is equivalent to the summed intensities of the daughters. The squared fluctuations in intensity of the two sibling cells is linearly related to the parent cell with a slope *α*, which is the fluorescence signal measured per partitioned repressor (right panel). Black points represent single divisions and red points are the means of 50 division events. Line corresponds to linear fit to the black points with a slope of *α* = 740 ± 40 a.u. per LacI.

While the mCherry fluorescence is proportional to the repressor copy number, it is not a direct measurement as the fluorescence of a single LacI-mCherry dimer is unknown *a priori*. Using video microscopy, we measure the partitioning statistics of the fluorescence intensity into two sibling cells after division [Fig. 2(C)]. This method, described in detail in the Materials and Methods and in Refs.(12, 25–27), reveals a linear relationship between the variance in intensity between two sibling cells and the intensity of the parent cell, the slope of which is equal to the brightness of a single LacI repressor. Since this measurement is performed simultaneously with measurement of the expression of the YFP reporter, this calibration factor was determined for each unique experimental replicate. We direct the reader to the SI text for a more thorough discussion of this inference.

### Scaling of gene expression with growth rate

Given the single-cell resolution of our experimental method, we examined how the cell volume and repressor copy number scaled across the different growth conditions at different levels of ATC induction. In agreement with the literature (1, 20, 28) our measurement reveals a strong linear dependence of the cell volume on the choice of carbon source, but no significant dependence on temperature [Fig. 3(A) and (B)]. Additionally, these findings are consistent across different ATC induction regimes. Together, these observations confirm that the particular details of our experimental system does not introduce unintended physiological consequences.

**Fig. 3.**
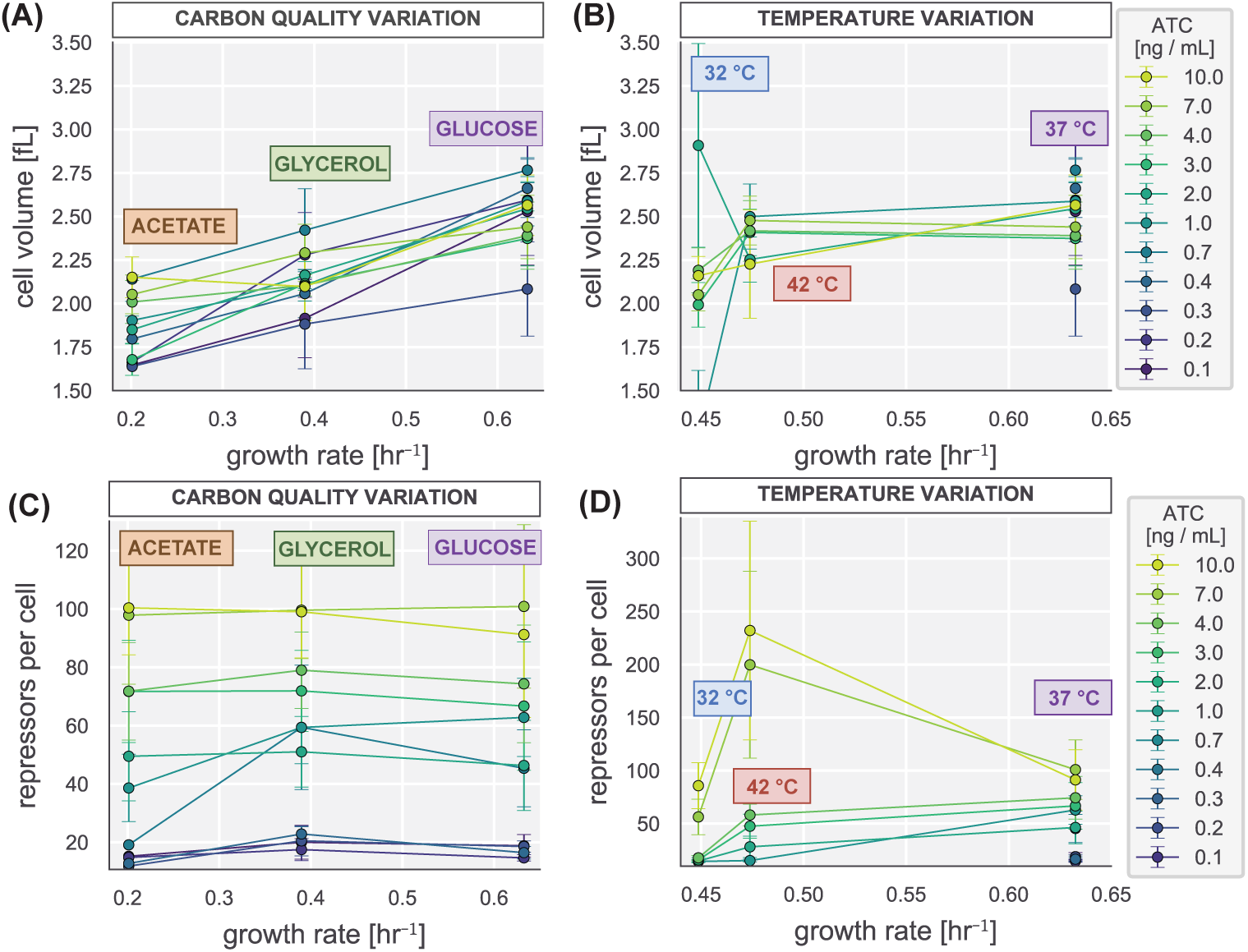
Scaling of cell size and repressor expression as a function of maximum growth rate. Dependence of cell volume on maximum growth rate under varying (A) carbon sources and (B) temperatures. Points and errors correspond to the mean and standard errors of five to eight biological replicates. The cell volume was calculated by approximating each cell as a spherocylinder and using measurements of the short and long axis lengths of each segmentation mask. Measured volumes are from snapshots of a nonsynchronously growing culture. The measured repressor copy number for each ATC induction condition (colored lines) as a function of the growth rate for various (C) carbon sources and (D) temperatures. Points and errors represent the mean and standard error of five to eight biological replicates. Colors correspond to the ATC induction concentration ranging from 10 ng/mL (yellow) to 0.1 ng/mL (black).

Using a fluorescence calibration factor determined for each experimental replicate [see Fig. 2(C) and Materials & Methods], we estimated the number of repressors per cell from snapshots of the mCherry signal intensity of each induction condition. Fig. 3(C) reveals a remarkable insensitivity of the repressor copy number on the growth rate under different carbon sources. Despite the change in cellular volume, the mean number of repressors expressed at a given induction condition is within error between all carbon sources. Previous work using mass spectrometry, a higher resolution method, has shown that there is a slight dependence of LacI copy number on growth rate expressed from its native promoter (5). It is possible that such a dependence exists in our experimental setup, but is not detectable with our lower resolution method. We also observe an insensitivity of copy number to growth rate when the temperature of the system is tuned [Fig. 3 (D)] though two aberrant points with large error obfuscates the presence of a growth rate dependence at high concentrations of ATC. For concentrations below 7 ng /mL, however, the repressor copy number remains constant across conditions. With no significant change in the repressor copy number and thus no dependence on the carbon source in our theoretical model, we are can immediately draw predictions of the fold-change in gene expression in different growth media.

### Fold-change dependence on carbon quality

Given *a priori* knowledge of the biophysical parameter values (10, 13) present in Eq. 2 and Eq. 3 and direct measurement of the repressor copy number, we made measurements of the fold-change in gene expression for each growth medium to test the prediction [Fig. 4 (A)]. We find that the measurements fall upon the predicted theoretical curve within error, suggesting that the values of the energetic terms in the model are insensitive to changing carbon sources. This is notable as glucose, glycerol, and acetate are metabolized via different pathways, changing the metabolite and protein composition of the cytosol (29, 30). This result underscores the utility of these thermodynamic parameters as quantitative traits in the study of growth-condition dependent gene expression.

**Fig. 4.**
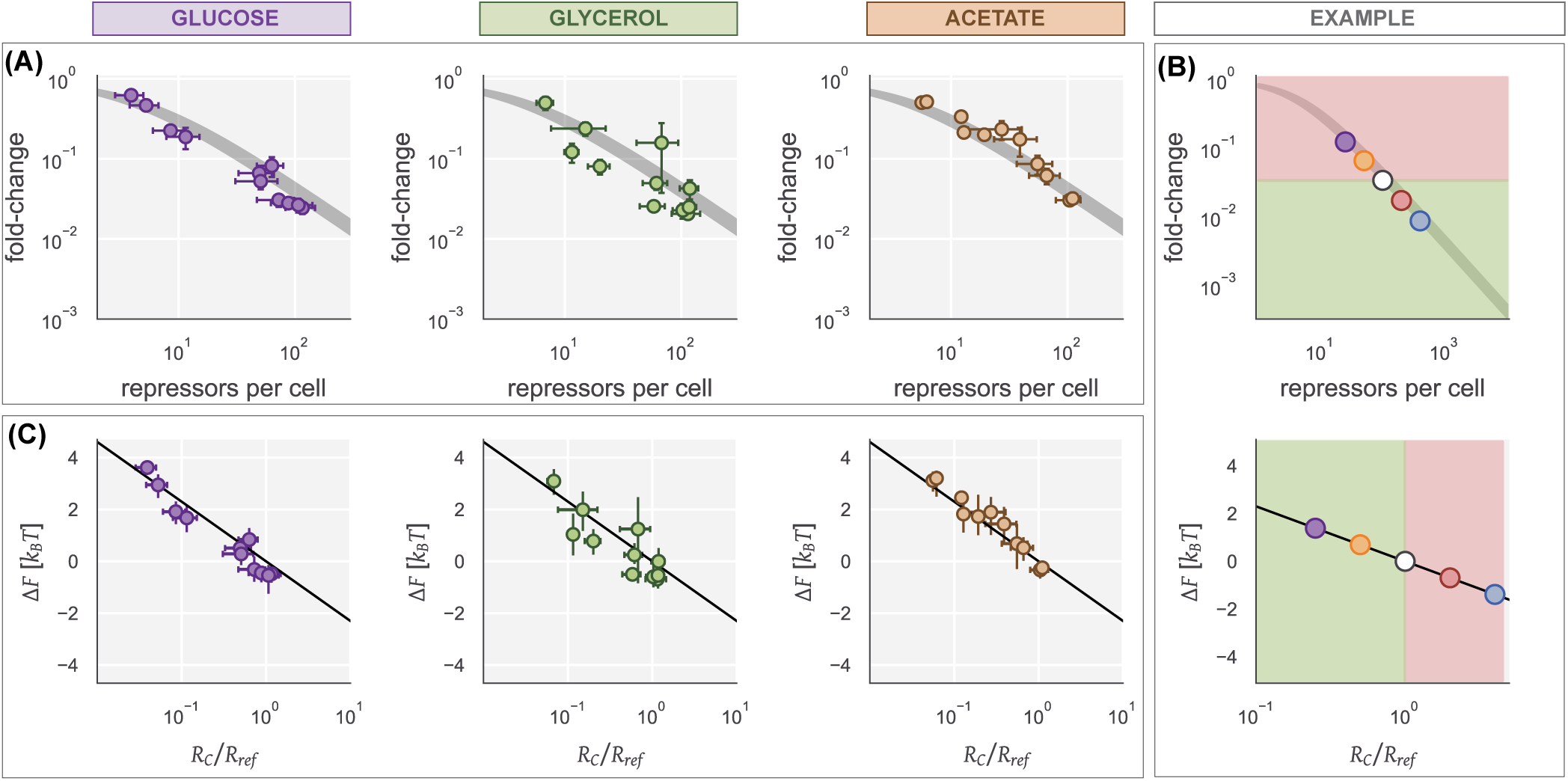
Fold-change in gene expression and free energy shifts in different growth media. (A) Measurements of the fold-change in gene expression plotted against the measured repressor copy number. Black line is the prediction and the width corresponds to the uncertainty in the DNA binding energy as reported in Ref. (10). Points and errors are the mean and standard error of five to eight replicates for each ATC induction condition. (B) An example of how fold-change is mapped onto changes in the free energy. Top panel shows the fold-change in gene expression at specific repressor copy numbers (points). Arbitrarily choosing 100 repressors per cell as the reference state (white point), any measurement with larger or smaller fold-change has a negative shift or positive shift in the free energy, shown as red and green panels respectively. If *R* is the only changing variable, the free energy shift should be a linear function of *R* (bottom panel). (C) The inferred free energy shift from the fold-change measurements in A. Black line is the prediction. Points correspond to the mean and standard error of the repressor copy number measurements over five to eight biological replicates. Vertical error bars correspond to the 95% credible region of the inferred free energy.

We have previously shown that Eq. 2 can be rewritten into a Fermi function of the form

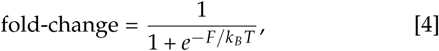

where *F* is the effective free energy difference between the repressor bound and unbound states of the promoter, also referred to as the Bohr parameter (9, 13, 31). For the case of an allosteric simple repression architecture, and given knowledge of the values of the biophysical parameters, *F* can be directly calculated as

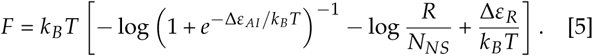

We have recently interrogated how this formalism can be used to map mutations within the repressor to biophysical parameters by examining the difference in free energy between a mutant and reference (wild-type) strain, Δ*F* = *F*_*mut*_ − *F*_*re f*_ (15). This approach revealed that different parametric changes yield characteristic response functions to changing inducer concentrations. Rather than using wild-type and mutant variants of the repressor, we can choose a reference condition and compare how the free energy changes between different growth media. Here, we choose the reference condition to be a sample grown at 37°C with glucose as the available carbon source and a repressor copy number *R* = 100 per cell. Under the hypothesis that the only variable parameter in these growth conditions is the repressor copy number *R*, the shift in free energy Δ*F* becomes

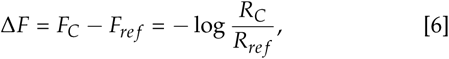

where *F*_*C*_ and *R*_*C*_ correspond to the free energy and repressor copy number of the different growth conditions. This concise prediction serves as a quantitative measure of how robust the energetic parameters Δ*ε*_*R*_ and Δ*ε* _*AI*_ are in units of *k*_*B*_*T* [Fig. 4(B)]. In using free energy shifts as a diagnostic, one can immediately determine the effect of the perturbation on the parameter values by quantifying the disagreement between the observed and predicted Δ*F* as the parameters and the free energy are both in the same natural units.

We inferred the observed free energy for the fold-change measurements shown in Fig. 4(A) [as described previously (15) and in the SI Text] and compared it to the theoretical prediction of Eq. 6, shown in Fig. 4(C). Again, we see the observed change in free energy is in strong agreement with our theoretical predictions. This agreement indicates that the free energy shift Δ*F* can be used in multiple contexts to capture the energetic consequences of physiological and evolutionary perturbations between different states of the system. The insensitivity of the biophysical parameters to these distinctly different physiological states demonstrates that Δ*ε* _*AI*_ and Δ*ε*_*R*_ are material properties of the repressor defined by the intricate hydrogen bonding networks of its constituent amino acids rather than by the chemical constituency of its surroundings. Tuning temperature, however, can change these material properties.

### Fold-change dependence on temperature variation

Unlike the identity of the carbon source, the temperature of the system is explicitly stated in Eq. 2 and Eq. 3 where Δ*ε*_*R*_ and Δ*ε* _*AI*_ are defined relative to the thermal energy of the system in which they were determined. This scaling is mathematically quantified as *k*_*B*_*T* dividing the exponentiated terms in Eq. 2 and Eq. 3. As all biophysical parameters were determined at a reference temperature of 37°C, any change in the growth temperature must be included as a correction factor. The simplest approach is to rescale the energy by the relative change in temperature. This is a simple multiplicative factor of *ϕ*_*T*_ = *k*_*B*_*T*_*re f*_ */k*_*B*_*T*_*exp*_where *T*_*re f*_ is the reference temperature of 37°C and *T*_*exp*_ is the experimental temperature. This is an intuitive result since an increase in temperature relative to the reference results in *ϕ*_*T*_ < 1, weakening the binding. Similarly, decreasing the temperature scales *ϕ*_*T*_ > 1, strengthening the binding relative to that of the reference temperature.

Fig. 5 (A) shows the measured fold-change in gene expression (points) plotted against the theoretical prediction with this correction factor (orange line). It is immediately evident that a simple rescaling of the energetic parameters is not sufficient for the 32°C condition and slightly underestimates the fold-change in the 42°C condition. To identify the source of this disagreement, we can again examine the free energy shift Δ*F*. As both Δ*ε* _*AI*_ and Δ*ε*_*R*_ are scaled to the thermal energy, Δ*F* defined as *F*_*T*_ − *F*_*re f*_ can be directly calculated as

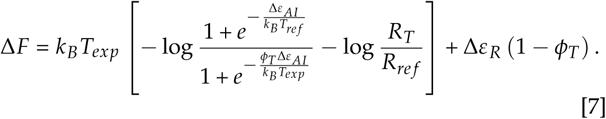

**Fig. 5.**
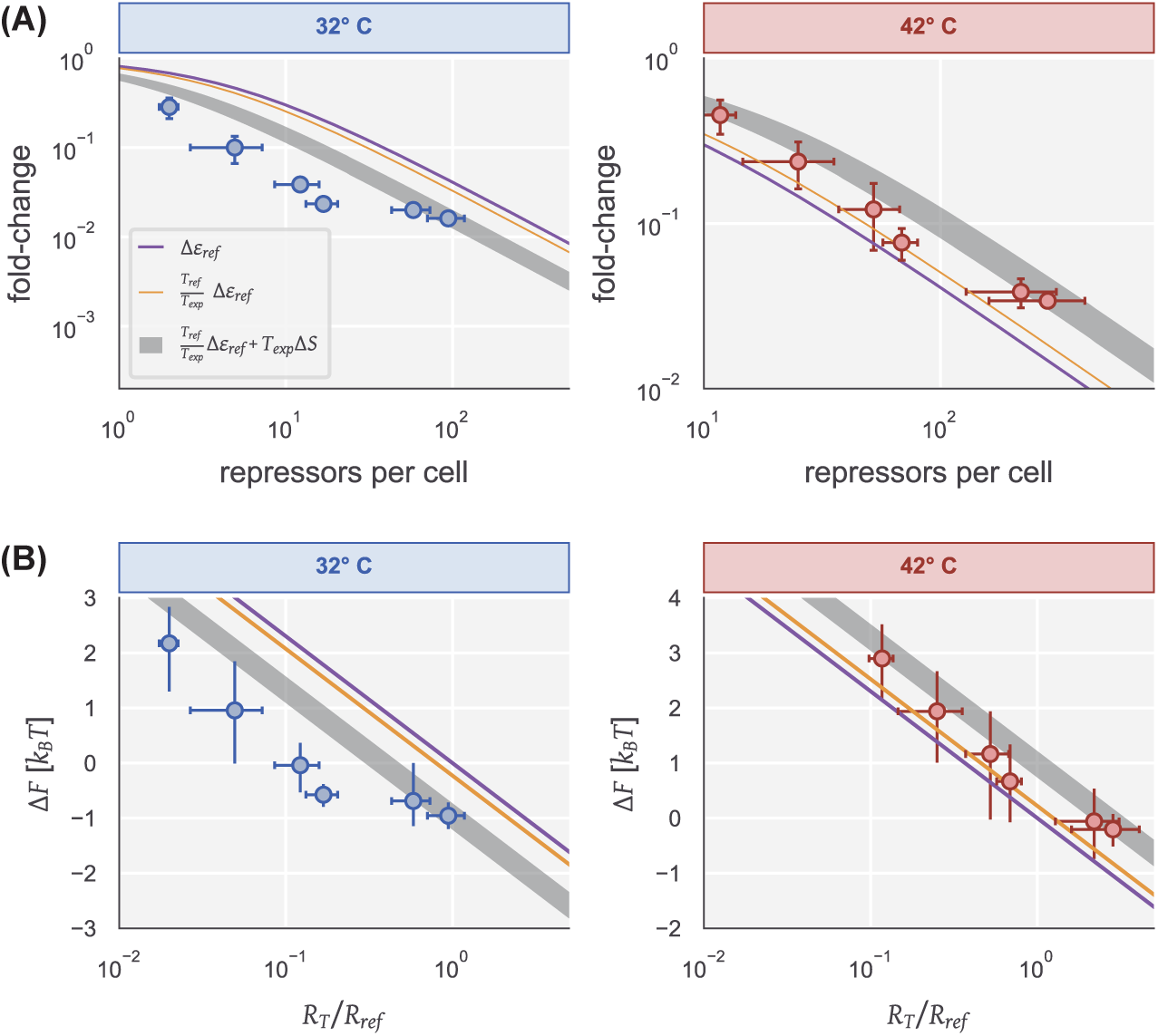
Temperature effects on the fold-change in gene expression and free energy. (A) The fold-change in gene expression for growth in glucose supplemented medium at 32°C (left) and 42°C (right). Points and errors correspond to the mean and standard error of five biological replicates. Predictions of the fold-change are shown without correcting for temperature (purple), with multiplicative scaling (orange), and with an entropic penalty (grey). The width of the prediction of the entropic penalty is the 95% credible region. (B) Predicted and observed shifts in free energy for growth glucose medium at 32°C (left) and 42°C (right). Points correspond to the median of the inferred shift in free energy. Vertical error bars indicate the bounds of the 95% credible region. Horizontal position and error corresponds to the mean and standard error for the repressor count over five biological replicates.

This prediction along with the empirically determined Δ*F* is shown in Fig. 5(B). Again, we see that this simple correction factor significantly undershoots or overshoots the observed Δ*F* for 32°C and 42°C, respectively, indicating that there are temperature dependent effects that are not accounted for in the simplest null model of temperature dependence.

The model described by Eqs. 2 and 3 subsumes the myriad rich dynamical processes underlying protein binding and conformational changes into two effective energies, Δ*ε*_*R*_ and Δ*ε* _*AI*_. By no means is this done to undercut the importance of these details in transcriptional regulation. Rather, it reduces the degrees of freedom in this objectively complex system to the set of the details critical to particular conditions in which we want to draw predictions. All prior dissections of this thermodynamic model have been performed at a single temperature, abrogating the need to consider temperature dependent effects. As we now vary temperature, we must consider details that are swept into the effective energies.

The model presented here only considers entropy by enumerating the multiplicity of states in which the repressor can bind to the DNA nonspecifically, resulting in terms of the form 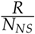. However, there are many other temperature-dependent entropic contributions to the effective energies such as the fraction of repressors bound to DNA versus in solution (32, 33), the vibrational entropy of the repressor (34), or conformational entropy of the genome (35, 36). We can consider the effective energies Δ*ε*_*R*_ and Δ*ε* _*AI*_ as having generic temperature dependent-entropic components Δ*S*_*R*_ and Δ*S*_*AI*_,

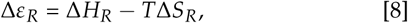

and

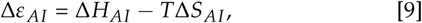

where Δ*H*_*R*_ and Δ*H*_*AI*_ is the enthalpic contribution to the energies Δ*ε*_*R*_ and Δ*ε* _*AI*_, respectively. Given the fold-change measurements at 32°C and 42°C, we estimated the entropic parameters Δ*S*_*R*_ and Δ*S*_*AI*_ under the constraints that at 37°C, Δ*ε*_*R*_ = −13.9 *k*_*B*_*T* and Δ*ε* _*AI*_ = 4.5 *k*_*B*_*T* (see the SI text for more discussion on this parameter estimation). The grey shaded lines in Fig. 5 show the result of this fit where the width represents the 95% credible region of the prediction given the estimated values of Δ*S*_*R*_ and Δ*S*_*AI*_. Including this phenomenological treatment of the entropy improves the prediction of the fold-change in gene expression [Fig. 5 (A)] as well as shift in free energy [Fig. 5 (B)]. This phenomenological description suggests that even small shifts in temperature can drastically alter the expression of a genetic circuit simply by tuning hidden entropic effects rather than scaling the difference in affinity between specific and nonspecific binding.

## DISCUSSION

The past century of work in bacterial physiology has revealed a rich molecular complexity that drives cellular growth rate (20). A key finding of this body of work is that the composition of the proteome is highly dependent on the specific growth condition, with entire classes of genes being up- or down-regulated to ensure that enough resources are allocated towards maintaining a pool of active ribosomes (2, 4). These studies have led to a coarse-grained view of global gene expression where physiological perturbations substantially change the molecular composition of the cell, obfuscating the utility of using thermodynamic models of individual regulatory elements across physiological states. In this work, we rigorously examine how robust the values of the various biophysical parameters are to changes in cellular physiology.

We first examined how nutrient fluctuations dictate the output of this architecture. We took three carbon sources with distinct metabolic pathways and varying quality and measured the level of gene expression, hypothesizing that the values of the biophysical parameters to be independent of the growth medium. We found that even when the growth rate is varied across a wide range (220 minute doubling time in acetate to 60 minute doubling time in glucose supported medium), there is no significant change to the fold-change in gene expression or in the expression of the transcription factor itself, within the resolution of our experiments. Given numerous quantitative studies of the proteomic composition reveal a dependence on protein content with growth rate (4–6), we find this robustness to be striking.

Schmidt *et al*. 2016 (5) found that the native expression of LacI has a weak positive correlation with the growth rate. The native LacI promoter region is solely regulated by activation via the cAMP Receptor Protein (CRP), a broadly acting dual regulator in *E. coli* (7). This is in contrast to the LacI expression system used in the present work where the promoter is negatively regulated by the TetR repressor, itself expressed from a low-copy number plasmid. Furthermore, the expression of LacI in this work is tuned by the addition of the allosteric effector of TetR, ATC, adding yet another layer of allosteric regulation on LacI expression. The significant difference in the regulatory mechanisms between the native and synthetic circuit used in this work makes the two findings difficult to directly compare. Regardless, our finding that the fold-change in gene expression is unaltered from one carbon source to another illustrates that the values of the biophysical parameters Δ*ε*_*R*_ and Δ*ε* _*AI*_ remain unperturbed, permitting quantitative prediction of gene expression across numerous physiological states.

However, in varying the temperature, we find that the predictive utility of the biophysical parameters values determined at 37°C is diminished, indicating that there are hidden effects not explicitly accounted for in our thermodynamic model. The measurements of the fold-change in gene expression are under- or over-estimated when the temperature is increased or decreased, respectively, when one simply rescales the energetic terms by the relative change in temperature. There are many features of transcriptional regulation that are not explicitly considered in our coarse-graining of the architecture into a two-state model. Recently, it has been suggested that the phenomenon of allostery writ large should be framed in the language of an ensemble of states rather than a simple active/inactive distinction (37). While our recent work illustrates that a two-state rendering of an allosteric repressor is highly predictive in a variety of situations (13, 15), we must now consider details which are dependent on the temperature of the system. In Fig. 5, we demonstrate that incorporating a temperature-dependent entropic cost to the energetic terms significantly improves the description of the experimental data. This is not to say, however, that this is now an open-and-closed case for what precisely defines this entropic cost. Rather, we conclude that the phenomenology of the temperature dependence can be better described by the inclusion of a correction factor that is linearly dependent on the system temperature. Biology is replete with phenomena which can introduce such an effect, including changes to the material properties of the repressor and DNA (34, 36), excluded volume effects (35), and solubilities (32, 33, 38). Understanding the mechanistic underpinnings of temperature dependence in elasticity theory was borne out of similar phenomenological characterization (39) and required a significant dialogue between theory and experiment (40). Further work is now needed to develop a theory of temperature effects in the regulation of gene expression.

The effective free energy *F*, as defined in Eq. 5, is a state variable of the simple repression regulatory architecture. This is illustrated in Fig. 6 where fold-change measurements from a wide array of conditions (and measurement techniques) can be collapsed onto the same theoretical description. Evolutionary perturbations (such as mutations in the operator or repressor sequence), physiological changes (such as modulations of the growth rate), or changes in the level of activity of the repressor (due to changes in inducer concentration) do not change the fundamental physics of the system and can all be described by changes in the free energy relative to one another. While such a statement is not “surprising”, we can now say it with quantitative confidence and use this principle to probe the degree to which physiological perturbations influence the biophysical parameters writ large. With such a framework in hand, we are in the auspicious position to take a predictive approach towards understanding how this regulatory architecture evolves in experimental settings, shedding light on the interplay between biophysical parameters, organismal fitness, and the fundamental forces of evolution.

**Fig. 6.**
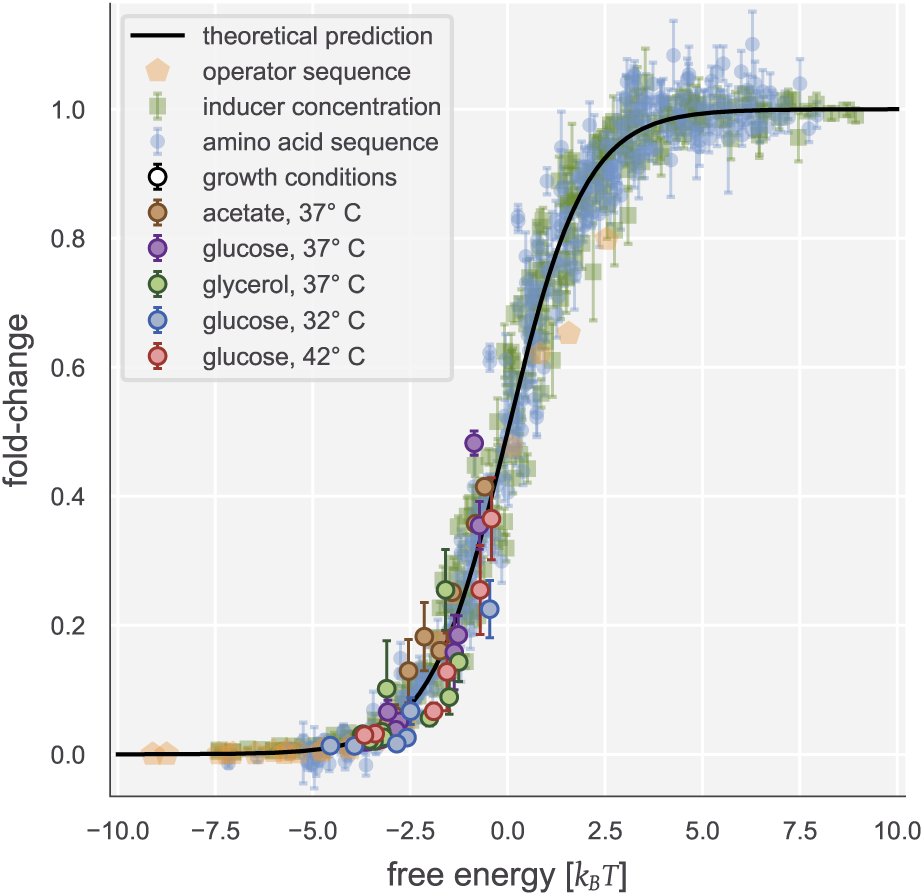
A singular theoretical description for the molecular biophysics of physiological and evolutionary adaptation in the simple repression regulatory architecture. Measurements of the fold-change in gene expression varying the sequence of the operator site [orange pentagons, Ref. (10)], concentration of extracellular inducer [green squares, Ref. (13)], amino-acid sequence of the repressor [blue points, Ref. (15)], and the various growth conditions queried in this work can be collapsed as a function of the effective free energy. Error bars correspond to the standard error of 5 to 10 biological replicates.

## Materials and Methods

### Bacterial Strains and Growth Media

Three genotypes were primarily used in this work, all in the genetic background of *Escherichia coli* MG1655-K12 and all derived from those used in Ref. (12). The three genotypes are as follows. For each experiment, an autofluorescence control was used which contained no fluorescent reporters [except for a CFP volume marker used for segmentation in Ref. (12)] which had the *lacI* and *lacZYA* genes deleted from the chromosome. The constitutive expression strain (Δ*lacI*; Δ*lacZYA*) included a YFP reporter gene integrated into the *galK* locus of the chromosome along with a kanamycin resistance cassette. The experimental strains in which LacI expression was controlled contained a *lacI-mCherry* fluorescent fusion integrated into the *ybcN* locus of the chromosome along with a chloramphenicol resistance cassette. This cassette was later deleted from the chromosome using FLP/FRT recombination (41, 42). The strain was then transformed with plasmid [pZS3-pN25-tetR following notation described in Ref. (43)] constitutively expressing the TetR repressor along with a chloramphenicol resistance cassette. All bacterial strains and plasmids used in this work are reported in the SI Text.

### Bacterial Growth Curves

Bacterial growth curves were measured in a multi-well plate reader (BioTek Cytation5) generously provided by the David Van Valen lab at Caltech. Cells constitutively expressing YFP were grown overnight to saturation in LB broth (BD Medical) at 37° C with aeration. Once saturated, cells were diluted 1000 fold into 50 mL of the desired growth medium and were allowed to grow at the appropriate experimental temperature with aeration for several hours until an OD_600*nm*_ ≈ 0.3 was reached. Cells were then diluted 1:10 into the desired growth media at the desired temperature. After being thoroughly mixed, 500 *µ*L aliquots were transferred to a black-walled 96-well plate (Brooks Automation Incorporated, Cat No. MGB096-1-2-LG-L), leaving two rows and two columns of wells on each side of the plate filled with sterile growth medium to serve as blanks and buffer against temperature variation. The plate was then transferred to the pre-warmed plate reader. OD_600*nm*_ measurements were made every five minutes for 12 to 24 hours until cultures had saturated. In between measurements, the plate incubated at the appropriate temperature with a linear shaking mode. We found that double-orbital shaking modes led to the formation of cell aggregates which gave inconsistent measurements.

### Estimation of Bacterial Growth Rate

Non-parametric estimation of the maximum growth rate was performed using the FitDeriv Python package as described in Ref. (44). Using this approach, the bacterial growth curve is modeled as a Gaussian process in which the measured growth at a given time point is modeled as a Gaussian distribution whose mean is dependent on the mean of the neighboring time points. This allows for smooth interpolation between adjacent measurements and calculation of second derivatives without an underlying parametric model. The reported growth rates are the maximum value inferred from the exponential phase of the experimental growth curve.

### Growth Conditions

Parent strains (autofluorescence control, Δ*lacI* constitutive control, and the ATC-inducible LacI-mCherry strain) were grown in LB Miller broth (B.D. Medical, Cat. No. 244620) at 37° C with vigorous aeration until saturated. Cells were then diluted between 1000 and 5000 fold into 3 mL of M9 minimal medium (B.D. Medical, Cat. No. 248510). The bacterial strain expressing the tetracycline-inducible LacI-mCherry was diluted into six 3 mL cultures with differing concentrations of ATC (Chemodex, Cat. No. CDX-A0197-T78) ranging from 0.1 to 10 ng / mL to induce expression of the transcription factor. These concentrations were reached by dilution from 1 *µ*g / mL stock in 50% ethanol. All cultures were shielded from ambient light using either aluminum foil or via an enclosure and were grown at the appropriate experimental temperature with aeration until an OD_600*nm*_ of approximately 0.25 − 0.35. Due to pipetting errors, cultures reached OD_600*nm*_ ≈ 0.3 at slightly different points in time. To ensure that strains could be directly compared, all strains were back diluted by several fold until equivalent. When all samples reached the appropriate OD_600*nm*_, the cells were harvested for imaging.

### Imaging Sample Preparation

Prior to the preparation of cell cultures for imaging, a 2% (w/v) agarose substrate (UltraPure, Thermo Scientific) was prepared and allowed to reach room temperature. For experiments conducted at 42°*C*, 4% (w/v) agarose substrates were prepared. Briefly, the agarose was mixed with the appropriate growth medium in a 50 mL conical polystyrene tube and then microwaved until molten. A 300 to 500 *µ*L aliquot was then sandwiched between two glass coverslips to ensure a flat imaging surface. Once solidified, the agarose pads were cut into squares approximately 0.5 cm per side.

To determine the calibration factor between fluorescence and protein copy number, production the fluorophore in question must be halted such that all differences in intensity between two daughter cells result from binomial partitioning of the fluorophores and not from continuing expression. This was achieved by removing the anhydrous tetracycline inducer from the growth medium through several washing steps as outlined in Ref. (12). Aliquots of 100 *µ*L from each ATC-induced culture were combined and pelleted at 13000×*g* for 2 minutes. The supernatant (containing ATC) was aspirated and replaced with 1 mL of M9 growth medium without ATC. The pellet was resuspended and pelleted at 13000 ×*g*. This wash step was repeated three times to ensure residual ATC had been removed and LacI-mCherry production was halted. After the final wash, the cell pellet was resuspended in 1 mL of M9 medium and diluted ten-fold. A 1 *µ*L aliquot of the diluted mixture was then spotted onto an agarose substrate containing the appropriate carbon source.

The remaining bacterial cultures (autofluorescence control, constitutive expression control, and the ATC-induced samples) were diluted ten-fold into sterile M9 medium. This dilution was thoroughly mixed and 1 *µ*L aliquots were spotted onto agarose substrates lacking the carbon source.

Once the spotted cells had dried onto the agarose substrates (about 5 to 10 minutes after deposition), the agarose pads were inverted and pressed onto a glass bottom dish (Electron Microscopy Sciences, Cat. No. 70674-52) and sealed with parafilm. Strips of double stick tape were added to the edge of the dish to help immobilize the sample on the microscope stage and minimize drift.

### Microscopy

All imaging was performed on a Nikon Ti-Eclipse inverted microscope outfitted with a SOLA LED fluorescence illumination system. All images were acquired on a Andor Zyla 5.5 sCMOS camera (Oxford Instruments Group). The microscope body and stage was enclosed in a plexiglass incubation chamber (Haison, approximately 1° C regulation control) connected to an external heater. Temperature of the stage was measured via a thermometer which controlled heating of the system.

All static images (i.e. images from which fold-change and repressor counts were calculated) were measured in an identical manner. Ten to fifteen fields of view containing on average 25 cells were imaged using phase contrast and fluorescence excitation. Fluorescence exposures were each 5 seconds while the phase contrast exposure time was between 75 ms and 150 ms. This procedure was repeated for each unique strain and ATC induction concentration.

To compute the calibration factor for that day of imaging, time-lapse images were taken of a separate agarose pad covered in cells containing various levels of LacI-mCherry. Fifteen to twenty positions were marked, choosing fields of view containing 20 to 50 cells. Cells were allowed to grow for a period of 90 to 120 minutes (depending on the medium-dependent growth rate) with phase contrast images taken every 5 to 10 minutes. At the final time-point, both phase contrast and fluorescence images were acquired using the same settings for the snapshots. Once the experiment was completed, images were exported to `.tif` format and transferred to cold storage and a computational cluster for analysis.

### Lineage Tracking

Cells were segmented and lineages reconstructed using the deep-learning-based bacterial segmentation software Super-Segger `v1.1` (45, 46) operated through MATLAB (Mathworks, `v2017b`). We found that the pre-trained network constants `100XEcM9` (packaged with the SuperSegger software) worked well for all growth conditions tested in this work. The generated files (`clist.mat`) associated with each sample and position were parsed using bespoke Python scripts to reconstruct lineages and apply appropriate filtering steps before calculating the fluorescence calibration factor. We direct the reader to the SI text for details of our data validation procedure to ensure proper lineage tracking.

### Calculation of the Calibration Factor

To estimate the calibration factor *α*, we used a Bayesian definition of probability to define a posterior distribution of *α* conditioned on intensity measurements of sibling cells after division. We direct the reader to the SI text for a detailed discussion of the inferential procedures and estimation of systematic error. We give a brief description of the inference below.

We are interested in determining the fluorescence of a single LacI-mCherry repressor dimer given a set of intensity measurements of sibling cells, [*I*_1_, *I*_2_]. The intensity of a given cell *I* is related to the number of LacI-mCherry dimers it is expressing by a multiplicative factor *α* which can be enumerated mathematically as

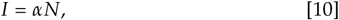

where *N* is the total number of LacI-mCherry dimers. We can define the posterior probability distribution of *α* conditioned on the intensity measurements using Bayes’ theorem as

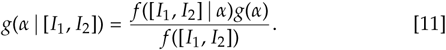

where we have used *g* and *f* as probability densities over parameters and data, respectively. The denominator of this expression (the evidence) is equivalent to the first term of the numerator (the likelihood) marginalized over *α*. In this work, this term serves as normalization factor and can be neglected.

Assuming that no more LacI-mCherry dimers are produced during cell division, the number of repressors that each sibling cell receives after division of the parent cell is binomially distributed with a probability *p*. We can make the approximation that partitioning of the repressors is even such that *p* = 1/2. The validity of this approximation is discussed in detail in the SI text. Using Eq. 10 and the change-of-variables formula, we can define the likelihood *g*([*I*_1_, *I*_2_] | *α*) as

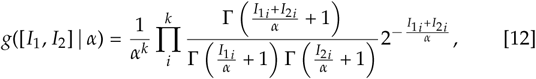

where *k* is the total number of sibling pairs measured.

With a likelihood defined, we must now define a functional form for *g*(*α*) which describes all prior information known about the calibration factor knowing nothing about the actual measurements. Knowing that we design the experiments such that only ≈ 2/3 of the dynamic range of the camera is used and *α* cannot be less than or equal to zero, we can define a half-normal distribution with a standard deviation of *σ* as

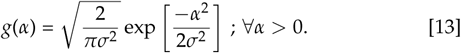

where the standard deviation is large, for example, *σ* = 500 a.u. / fluorophore. We evaluated the posterior distribution using Markov chain Monte Carlo (MCMC) as is implemented in the Stan probabilistic programming language (47). The `.stan` file associated with this model along with the Python code used to execute it can be accessed on the paper website.

### Counting Repressors

Given an estimation for *α* for each experiment, we calculate the total number of repressors per cell from a

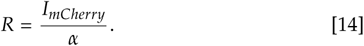

However, as discussed in detail in the SI text, a systematic error in the repressor count is introduced due to division in the asynchronous culture between the cessation of LacI-mCherry production and the actual imaging. The entire sample preparation procedure is ≈30 − 60 min, during which time some cells complete a cell division, thereby diluting the total repressor count. To ensure that the measured number of repressors corresponded to the measured YFP intensity, we restricted our dataset for all experiments to cells that had a pole-to-pole length *ℓ* ≥ 3.5 *µ*m, indicating that they had likely not undergone a division during the sample preparation.

### Code and Data Availability

All code used in this work is available on the paper website and associated GitHub repository(DOI: 0.5281/zenodo.3560369). This work also used the open-source software tools SuperSegger `v.1.1`(45, 46) for lineage tracking and FitDeriv `v.1.03` for the nonparametric estimation of growth rates. Raw image files are large (≈ 1.8 Tb) and are therefore available upon request. The clist.mat files used to calculate fold-change and to assign sibling cells can be accessed via the associated GitHub repository via (DOI: 0.5281/zenodo.3560369) or through Caltech DATA under the DOI: 10.22002/D1.1315.

## ACKNOWLEDGMENTS

We thank Prof. David Van Valen, Ed Pao, Geneva Miller, and Andy Halleran for access and training for the use of the Biotek plate reader for growth rate measurements. The experimental efforts of this work first took place during the Physiology course at the Marine Biological Laboratory operated by the University of Chicago in Woods Hole, MA, USA. We thank Celine Akelmade, George Bell, Cayla Jewett, Emily Meltzer, and Elizabeth Mueller for their work on the project during the course. We also thank Suzannah Beeler, Nathan Belliveau, Justin Bois, Rob Brewster, Soichi Hirokawa, Heun Jin Lee, Muir Morrison, Manuel Razo-Mejia, and Gabe Salmon for extensive advice and discussion. This work was supported by NIH funding under 1R35 GM118043 Maximizing Investigators’ Research Award (MIRA) and through the John Templeton Foundation as part of the Boundaries of Life Initiative via grants 51250 and 60973.

The authors declare no conflict of interest.

## Notes

https://rpgroup.caltech.edu/mwc_growth

https://github.com/rpgroup-pboc/mwc_growth

## References

1. Schaechter M, Maaløe O, Kjeldgaard NO (1958) Dependency on Medium and Temperature of Cell Size and Chemical Composition during Balanced Growth of *salmonella Typhimurium*. Microbiology 19(3):592–606.

2. Scott M, Klumpp S, Mateescu EM, Hwa T (2014) Emergence of robust growth laws from optimal regulation of ribosome synthesis. Molecular Systems Biology 10(8):747–747.

3. Klumpp S, Hwa T (2014) Bacterial growth: Global effects on gene expression, growth feed-back and proteome partition. Current Opinion in Biotechnology 28:96–102.

4. Hui S, et al. (2015) Quantitative proteomic analysis reveals a simple strategy of global resource allocation in bacteria. Molecular Systems Biology 11(2).

5. Schmidt A, et al. (2016) The quantitative and condition-dependent *Escherichia* coli proteome. Nature Biotechnology 34(1):104–110.

6. Li GW, Burkhardt D, Gross C, Weissman JS (2014) Quantifying Absolute Protein Synthesis Rates Reveals Principles Underlying Allocation of Cellular Resources. Cell 157(3):624–635.

7. Gama-Castro S, et al. (2016) RegulonDB version 9.0: High-level integration of gene regulation, coexpression, motif clustering and beyond. Nucleic Acids Research 44(D1):D133–D143.

8. Rydenfelt M, Garcia HG, Cox RS, Phillips R (2014) The Influence of Promoter Architectures and Regulatory Motifs on Gene Expression in *Escherichia coli*. PLoS ONE 9(12):e114347.

9. Phillips R, et al. (2019) Figure 1 Theory Meets Figure 2 Experiments in the Study of Gene Expression. Annual Review of Biophysics 48(1):121–163.

10. Garcia HG, Phillips R (2011) Quantitative dissection of the simple repression input-output function. Proceedings of the National Academy of Sciences 108(29):12173–12178.

11. Garcia HG, et al. (2012) Operator Sequence Alters Gene Expression Independently of Transcription Factor Occupancy in Bacteria. Cell Reports 2(1):150–161.

12. Brewster RC, et al. (2014) The Transcription Factor Titration Effect Dictates Level of Gene Expression. Cell 156(6):1312–1323.

13. Razo-Mejia M, et al. (2018) Tuning Transcriptional Regulation through Signaling: A Predictive Theory of Allosteric Induction. Cell Systems 6(4):456–469.e10.

14. Barnes SL, Belliveau NM, Ireland WT, Kinney JB, Phillips R (2019) Mapping DNA sequence to transcription factor binding energy in vivo. PLOS Computational Biology 15(2):e1006226.

15. Chure G, et al. (2019) Predictive Shifts in Free Energy Couple Mutations to Their Phenotypic Consequences. Proceedings of the National Academy of Sciences 116(37).

16. Jones DL, Brewster RC, Phillips R (2014) Promoter architecture dictates cell-to-cell variability in gene expression. Science 346(6216):1533–1536.

17. Ko MSH (1991) A stochastic model for gene induction. Journal of Theoretical Biology 153(2):181–194.

18. Kepler TB, Elston TC (2001) Stochasticity in Transcriptional Regulation: Origins, Consequences, and Mathematical Representations. Biophysical Journal 81(6):3116–3136.

19. Michel D (2010) How transcription factors can adjust the gene expression floodgates. Progress in Biophysics and Molecular Biology 102(1):16–37.

20. Jun S, Si F, Pugatch R, Scott M (2018) Fundamental principles in bacterial physiology—history, recent progress, and the future with focus on cell size control: A review. Reports on Progress in Physics 81(5):056601.

21. Weinert FM, Brewster RC, Rydenfelt M, Phillips R, Kegel WK (2014) Scaling of Gene Expression with Transcription-Factor Fugacity. Physical Review Letters 113(25).

22. Shahab N, Flett F, Oliver SG, Butler PR (1996) Growth rate control of protein and nucleic acid content in *streptomyces coelicolor* A3(2) and *escherichia coli* B/r. Microbiology, 142(8):1927–1935.

23. Zaritsky A, Woldringh CL (1978) Chromosome replication rate and cell shape in *Escherichia* coli: Lack of coupling. Journal of Bacteriology 135(2):581–587.

24. Liu M, et al. (2005) Global Transcriptional Programs Reveal a Carbon Source Foraging Strategy by *Escherichia coli*. Journal of Biological Chemistry 280(16):15921–15927.

25. Rosenfeld N, Young JW, Alon U, Swain PS, Elowitz MB (2005) Gene Regulation at the Single-Cell Level. Science 307(5717):1962–1965.

26. Rosenfeld N, Perkins TJ, Alon U, Elowitz MB, Swain PS (2006) A Fluctuation Method to Quantify In Vivo Fluorescence Data. Biophysical Journal 91(2):759–766.

27. Teng SW, et al. (2010) Measurement of the Copy Number of the Master Quorum-Sensing Regulator of a Bacterial Cell. Biophysical Journal 98(9):2024–2031.

28. Shehata TE, Marr AG (1975) Effect of temperature on the size of *Escherichia* coli cells. Journal of Bacteriology 124(2):857–862.

29. Martínez-Gómez K, et al. (2012) New insights into *Escherichia* coli metabolism: Carbon scavenging, acetate metabolism and carbon recycling responses during growth on glycerol. Microbial Cell Factories 11:46.

30. Kim PJ, et al. (2007) Metabolite essentiality elucidates robustness of *Escherichia* coli metabolism. Proceedings of the National Academy of Sciences of the United States of America 104(34):13638–13642.

31. Phillips R (2015) Napoleon Is in Equilibrium. Annual Review of Condensed Matter Physics 6(1):85–111.

32. Elf J, Li GW, Xie XS (2007) Probing Transcription Factor Dynamics at the Single-Molecule Level in a Living Cell. Science 316(5828):1191–1194.

33. Kao-Huang Y, et al. (1977) Nonspecific DNA binding of genome-regulating proteins as a biological control mechanism: Measurement of DNA-bound *Escherichia* coli lac repressor in vivo. Proceedings of the National Academy of Sciences 74(10):4228–4232.

34. Goethe M, Fita I, Rubi JM (2015) Vibrational Entropy of a Protein: Large Differences between Distinct Conformations. Journal of Chemical Theory and Computation 11(1):351–359.

35. Driessen RPC, et al. (2014) Effect of Temperature on the Intrinsic Flexibility of DNA and Its Interaction with Architectural Proteins. Biochemistry 53(41):6430–6438.

36. Mondal J, Bratton BP, Li Y, Yethiraj A, Weisshaar JC (2011) Entropy-Based Mechanism of Ribosome-Nucleoid Segregation in E. coli Cells. Biophysical Journal 100(11):2605–2613.

37. Motlagh HN, Wrabl JO, Li J, Hilser VJ (2014) The ensemble nature of allostery. Nature 508(7496):331–339.

38. Yakovchuk P, Protozanova E, Frank-Kamenetskii MD (2006) Base-stacking and base-pairing contributions into thermal stability of the DNA double helix. Nucleic Acids Research 34(2):564–574.

39. Friedel J (1974) On the stability of the body centred cubic phase in metals at high temperatures. Journal de Physique Lettres 35(4):59–63.

40. Phillips R (2001) Crystals, Defects and Microstructures by Rob Phillips. (Cambridge Publishing).

41. Schlake T, Bode J (1994) Use of Mutated FLP Recognition Target (FRT) Sites for the Exchange of Expression Cassettes at Defined Chromosomal Loci. Biochemistry 33(43):12746–12751.

42. Zhu XD, Sadowski PD (1995) Cleavage-dependent Ligation by the FLP Recombinase Characterization of a Mutant FLP Protein With An Alteration In a Catalytic Amino Acid. Journal of Biological Chemistry 270(39):23044–23054.

43. Lutz R, Bujard H (1997) Independent and Tight Regulation of Transcriptional Units in *Escherichia coli* Via the LacR/O, the TetR/O and AraC/I1-I2 Regulatory Elements. Nucleic Acids Research 25(6):1203–1210.

44. Swain PS, et al. (2016) Inferring time derivatives including cell growth rates using Gaussian processes. Nature Communications 7:13766.

45. Stylianidou S, Brennan C, Nissen SB, Kuwada NJ, Wiggins PA (2016) SuperSegger: Robust image segmentation, analysis and lineage tracking of bacterial cells. Molecular Microbiology 102(4):690–700.

46. Cass JA, Stylianidou S, Kuwada NJ, Traxler B, Wiggins PA (2017) Probing bacterial cell biology using image cytometry. Molecular Microbiology 103(5):818–828.

47. Carpenter B, et al. (2017) Stan: A Probabilistic Programming Language. Journal of Statistical Software 76(1):1–32.

