## Supplemental Information for "Physiological Adaptability and Parametric Versatility in a Simple Genetic Circuit"

**8 Media Recipes and Bacterial Strains** **23**

### 1. Derivation of the Simple Repression Input-Output Function

Here, we derive the functional form of the thermodynamic model given as Eqns. 2 and 3 in the main text of this work. We direct the interested reader to Refs. (1, 2) for a more thorough discussion of the derivation and exploration of the properties of the input-output function.

We begin by defining an inducible simple repression architecture as a regulatory motif in which the binding of a transcription factor (namely, a repressor) to a specific site on the genome occludes binding of an RNA polymerase (RNAP), thereby preventing transcription of the target gene [Fig. S1(A)]. Here, we declare the repressor to be an allosteric molecule, meaning that it fluctuates between active and inactive states in thermal equilibrium. The proportion of the repressor pool in the inactive state is governed by the binding of a small molecule called an inducer to an auxiliary site on the repressor. When an inducer is bound, the inactive state becomes energetically preferable and transcription can proceed. Both states of the repressor are capable of binding the DNA, but the inactive state binds with significantly weaker affinity than the active state.

Ultimately, we wish to compute the level of gene expression as a function of the various affinities and copy numbers of the regulatory components. We make the approximation that the level of gene expression is proportional to the probability of an RNAP being bound to the promoter,

$$\textit{gene-expression} \propto P_{bound}. \tag{1}$$

To compute this probability, we make the approximation that on the time-scale at which binding and unbinding of molecular species occurs, the system is in a quasi-equilibrium. This allows us to say that the probability of a polymerase or repressor being bound to its cognate binding site obeys a Boltzmann distribution

$$P_{state} \propto e^{-\varepsilon_{state}/k_B T}, \tag{2}$$

where $\varepsilon_{state}$ is the energy of that particular state, $k_B$ is the Boltzmann constant, and $T$ is the temperature of the system in Kelvin. With this in hand, we can enumerate all of the occupancy states of the promoter, shown in Fig. S1(B), normalizing the statistical weight to the unoccupied promoter. Here, we use $P$, $R_A$, and $R_I$ to denote the average copy number of the polymerase, active repressors, and inactive repressors, respectively. The multiplicity of states in which they can be bound to the DNA can be approximated by dividing the copy number by the number of nonspecific sites $N_{NS}$. As there is only one specific site $N_S$ in the genome, $N_{NS} \gg N_S$ and thus we can approximate $N_{NS}$ as the length of the *E. coli* genome in base pairs, $\approx 5 \times 10^6$. The free energy difference between nonspecific and specific binding of the polymerase, active repressor, and inactive repressor are denoted as $\Delta\varepsilon_P$, $\Delta\varepsilon_{RA}$, and $\Delta\varepsilon_{RI}$, respectively.

Given the enumerated states and weights, we can calculate the probability of the promoter being occupied by an RNA polymerase $P_{bound}$ directly as

$$P_{bound} = \frac{\frac{P}{N_{NS}} e^{-\beta\Delta\varepsilon_P}}{1 + \frac{P}{N_{NS}} e^{-\beta\Delta\varepsilon_P} + \frac{R_A}{N_{NS}} e^{-\beta\Delta\varepsilon_{RA}} + \frac{R_I}{N_{NS}} e^{-\beta\Delta\varepsilon_{RI}}}, \tag{3}$$

where we have used the shorthand notation of $\beta = \frac{1}{k_B T}$.

Experimental measurement of $P_{bound}$ is fraught with difficulty because determination of the proportionality to gene expression is not straightforward. However, we can compute the fold-change in gene expression, defined as the level of gene expression in the presence of repressors relative to a constitutive promoter, resulting in

$$\textit{fold-change} = \frac{P_{bound}(R_A + R_I > 0)}{P_{bound}(R_A + R_I = 0)} = \frac{1 + \frac{P}{N_{NS}} e^{-\beta\Delta\varepsilon_P}}{1 + \frac{P}{N_{NS}} e^{-\beta\Delta\varepsilon_P} + \frac{R_A}{N_{NS}} e^{-\beta\Delta\varepsilon_{RA}} + \frac{R_I}{N_{NS}} e^{-\beta\Delta\varepsilon_{RI}}}. \tag{4}$$

This result can be further simplified by making two assumptions. First, we assume that binding of the inactive repressor to the promoter is comparable to nonspecific binding, meaning that $\frac{R_I}{N_{NS}} e^{-\beta\Delta\varepsilon_{RI}} \ll 1 + \frac{R_A}{N_{NS}} e^{-\beta\Delta\varepsilon_{RA}}$ and can therefore be neglected. Additionally, we make the approximation that the promoter is weak, meaning that $\frac{P}{N_{NS}} e^{-\beta\Delta\varepsilon_P} \ll 1$. This assumption is not unreasonable as the average polymerase copy number is on the order of $\approx 10^3$ per cell (3) and characteristic polymerase-DNA binding energies are $\approx 2 - 5\,k_B T$ (4). Together, these estimates bring the quantity $\frac{P}{N_{NS}} e^{-\beta\Delta\varepsilon_P} \approx 0.001$, which we consider to be a negligible probability. Under these assumptions, Eq. 4 can be simplified to a tidy form of

$$\textit{fold-change} = \frac{1}{1 + \frac{R_A}{N_{NS}} e^{-\beta\Delta\varepsilon_{RA}}}. \tag{5}$$

We are now tasked with determining the average copy number of *active* repressors per cell, a quantity that evades direct *in vivo* experimental measurement. By modeling the repressor as an allosteric molecule, we can compute probability of a repressor being in the active state $P_{act}$ by again considering the states and statistical weights of the system, illustrated in Fig. S1(C). Here, we've used $c$ to denote the concentration of extracellular inducer, $K_A$ and $K_I$ to denote the dissociation constants of the inducer to the active and inactive repressor, respectively, and $\Delta\varepsilon_{AI}$ to denote the energy difference between the two states. The quantity $p_{act}$ can be be calculated as

$$p_{act}(c) = \frac{\left(1 + \frac{c}{K_A}\right)^2}{\left(1 + \frac{c}{K_A}\right)^2 + e^{-\beta\Delta\varepsilon_{AI}}\left(1 + \frac{c}{K_I}\right)^2}, \tag{6}$$

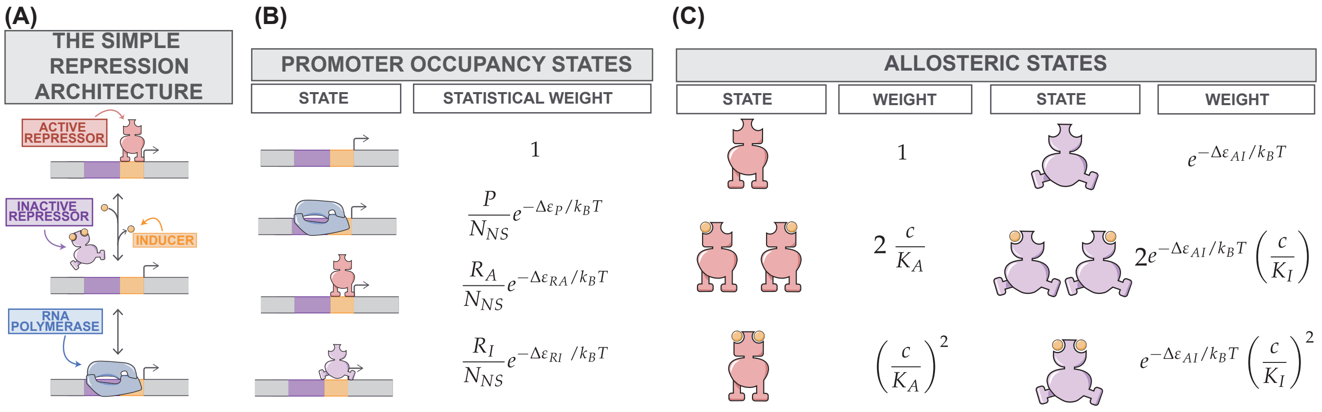

**Fig. S1. States and statistical weights for the simple repression architecture.** (A) A schematic of the inducible simple repression architecture. An active repressor (red) binds preferentially to its cognate binding site (orange rectangle), occluding binding of an RNA polymerase (blue). In the presence of a small molecule inducer (orange sphere), the inactive state of the repressor (purple) becomes more energetically favorable than the active state. This results in loss of specific binding to the cognate site, which in turn permits binding of RNA polymerase to the promoter (purple rectangle) and transcription to occur. (B) States and statistical weights for occupancy of the promoter. All weights are computed relative to the unoccupied promoter state (top). $R_A$ and $R_I$ correspond to the total number of active and inactive repressors present in the cell, respectively. The energies $\Delta\varepsilon_{RA}$ and $\Delta\varepsilon_{RI}$ are the repressor-DNA binding energies for the active and inactive states, respectively. $P$ is the the total number of RNA polymerases per cell and $\Delta\varepsilon_P$ is its binding energy to the promoter sequence. (C) Allosteric states and statistical weights of the repressor. All weights are computed relative to the active state in the absence of inducer. Variables $K_A$ and $K_I$ correspond to the dissociation constants for the active and inactive states of the repressor to the inducer molecule, respectively. Concentration of the inducer is denoted as $c$. The energetic parameter $\Delta\varepsilon_{AI}$ is the energetic difference between the active and inactive states.

where the factor of 2 in the superscript illustrates that the repressor has two inducer binding sites.

Using Eq. 6, we can now write the expression for the fold-change in Eq. 5 in terms of the *total* repressor copy number $R = R_A + R_I$ as

$$\textit{fold-change} = \frac{1}{1 + p_{act}(c)\frac{R}{N_{NS}}e^{-\beta\Delta\varepsilon_{RA}}}. \qquad [7]$$

In this work, we only consider the fold-change in gene expression in the absence of inducer ($c = 0$), abrogating the need to consider the inducer dissociation constants. In this particular condition, Eq. 6 can be simplified further as

$$p_{act}(c = 0) = \frac{1}{1 + e^{-\beta\Delta\varepsilon_{AI}}}, \qquad [8]$$

which is equivalent to Eq. 3 in the main text. Putting together Eq. 8 and Eq. 7 yields the expression

$$\textit{fold-change} = \frac{1}{1 + p_{act}(c = 0)\frac{R}{N_{NS}}e^{-\beta\Delta\varepsilon_{RA}}}, \qquad [9]$$

which is the same as Eq. 2 in the main text and the centerpiece of this work. In the main text of this work, $\Delta\varepsilon_R$ was used in lieu of $\Delta\varepsilon_{RA}$ for notational clarity.

### 2. Non-parametric Inference of Growth Rates

In this section, we discuss the measurement of the bacterial growth curves as well as our strategy for estimating the growth rates for all experimental conditions in this work.

**A. Experimental growth curves.** As is described in the Materials and Methods section of the main text, we measured the growth of *E. coli* strains constitutively expressing YFP using a BioTek Cytation 5 96-well plate reader generously provided by the lab of Prof. David Van Valen at Caltech. Briefly, cells were grown overnight in a nutrient rich LB medium to saturation and were subsequently diluted 1000 fold into the appropriate growth medium. Once these diluted cultures reached an $OD_{600nm}$ of $\approx 0.3$, the cells were again diluted 100 fold into fresh medium preheated to the appropriate temperature . Aliquots of 300 $\mu$L of this dilution were then transferred to a 96-well plate leaving two-rows on all sides of the plate filled with sterile media to serve as blank measurements. Once prepared, the plate was transferred to the plate reader and $OD_{600nm}$ measurements were measured every $\approx$ 5 - 10 min for 12 to 18 hours. Between measurements, the plates were agitated with a linear shaking mode to avoid sedimentation of the culture. A series of technical replicates of the growth curve in glycerol supplemented medium at 37° C is shown in Fig. S2(A).

**B. Inference of maximum growth rate.** The phenomenon of collective bacterial growth has been the subject of intense research for the better part of a century (5, 6) yielding many parametric descriptions of the bulk growth rates each with varying degrees of detail (6, 7). For this scope of this work, we are not particularly interested in estimating numerous parameters that describe the phenomenology of the growth curves. Rather, we are interested in knowing a single quantity – the maximum growth rate – and the degree to which it is tuned across the different experimental conditions. To avoid forcing the bacterial growth curves into a parametric form, we treated the observed growth curves using Gaussian process modeling using as implemented in the FitDeriv software described in Ref. (8). This method permits an estimation of the most-likely $OD_{600nm}$ value at each point in time given knowledge of the adjacent measurements. The weighting given to all points in the series of measurements is defined by the covariance kernel function and we direct the reader to Ref. (8) for a more detailed discussion on this kernel choice and overall implementation of Gaussian process modeling of time-series data. As this approach estimates the probability of a given $OD_{600nm}$ at each time point, we can compute the mean and standard deviation. Fig. S2(B) show the raw measurements and the mean estimated value in green and purple, respectively. With the high temporal resolution of the $OD_{600nm}$ measurements , modeling the entire growth curve becomes a computationally intensive task. Furthermore, as we are interested in only the maximum growth rate, there is no benefit in analyzing the latter portion of the experiment where growth slows and the population reaches saturation. We therefore manually restricted each analyzed growth curve to a region capturing early and mid exponential phase growth, illustrated by the shaded region in Fig. S2(A).

With a smooth description of the $OD_{600nm}$ measurements as a function of time with an appropriate measure of uncertainty, we can easily compute the time derivative which is equivalent to the growth rate. A representative time derivative is shown in Fig. S2(C). Here, the dark purple curve is the mean value of the time derivative and the shaded region is $\pm$ one standard deviation. The maximum value of this inferred derivative is the reported maximum growth rate of that experimental condition.

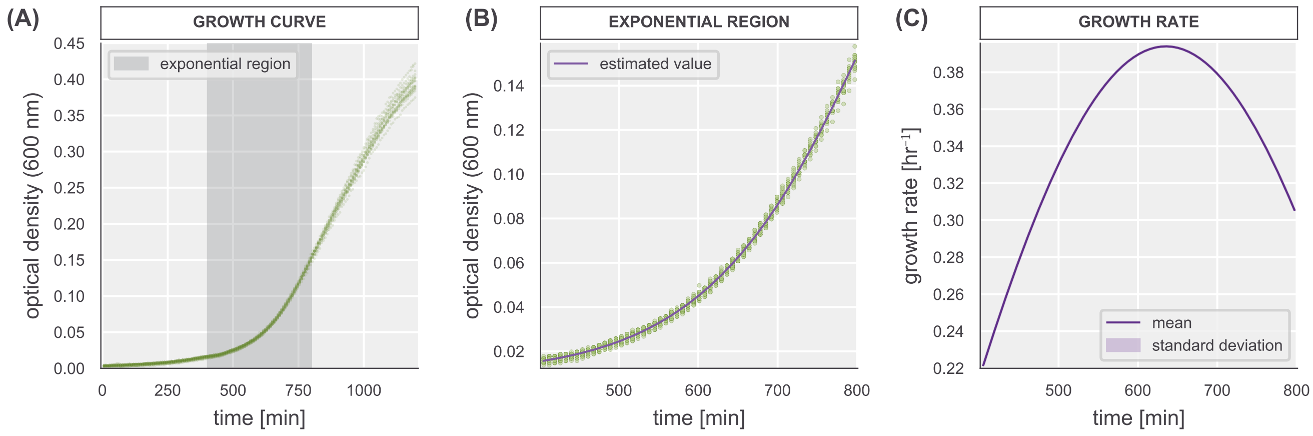

**Fig. S2. Non-parametric characterization of bacterial growth curves and estimation of the growth rate.** (A) Representative biological replicate growth curve of a $\Delta lacIZYA$ *E. coli* strain in glycerol supplemented M9 minimal media at 37° C. Points represent the individual optical density measurements across eight technical replicates. The shaded region illustrates the time window from which the maximum growth rate was inferred. (B) Green points are those from the shaded region in (A). Purple line is the estimated value of the optical density resulting from the gaussian process modeling. (C) The value of the first derivative of the optical density as a function of time estimated via Gaussian process modeling. The first derivative is taken as the growth rate at each time point. The solid line is the estimated mean value and the shaded region represents the standard deviation of the posterior distribution as calculated by the `FitDeriv` software (8).

### 3. Approximating Cell Volume

In Fig. 2 of the main text, we make reference to the volume of the cells grown in various conditions. Here, we illustrate how we approximated this estimate using measurements of the individual cell segmentation masks.

Estimation of bacterial cell volume and its dependence on the total growth rate has been the target of numerous quantitative studies using a variety of methods including microscopy (5, 9–12) and microfluidics (13) revealing fascinating phenomenology of growth at the single cell level . Despite the high precision and extensive calibration of these methods, it is not uncommon to have different methods yield different estimates, indicating that it is not a trivial measurement to make. In the present work, we sought to estimate the cell volume and compare it to the well-established empirical results of the field of bacterial physiology to ensure that our experimental protocol does not alter the physiology beyond expectations. As the bulk of this work is performed using single-cell microscopy, we chose to infer the approximate the cell volume from the segmentation masks produced by the SuperSegger MATLAB software (14) which reported the cell length and width in units of pixels which can be converted to meaningful units given knowledge of the camera interpixel distance.

We approximated each segmented cell as a cylinder of length $a$ and radius $r$ capped on each end by hemispheres with a radius $r$. With these measurements in hand, the total cell volume was computed as

$$V_{cell} = \pi r^2 \left(a + \frac{4}{3}r\right). \quad [10]$$

The output of the SuperSegger segmentation process is an individual matrix for each cell with a variety of fluorescence statistics and information regarding the cell shape. Of the latter category, the software reports in pixels the total length $\ell$ and width $w$ of the cell segmentation mask. Given these measurements, we computed the radius $r$ of the spherocylinder as

$$r = \frac{w}{2} \quad [11]$$

and the cylinder length $a$ as

$$a = \ell - w. \quad [12]$$

Fig. S3 shows the validity of modeling the segmentation masks as a spherocylinder in two dimensions. Here, the thin colored lines are the contours of a collection of segmentation masks and the black solid lines are spherocylinders using the average length and width of the segmentation masks, as calculated by 11 and Eq. 12. It appears that this simple approximation is reasonable for the purposes of this work.

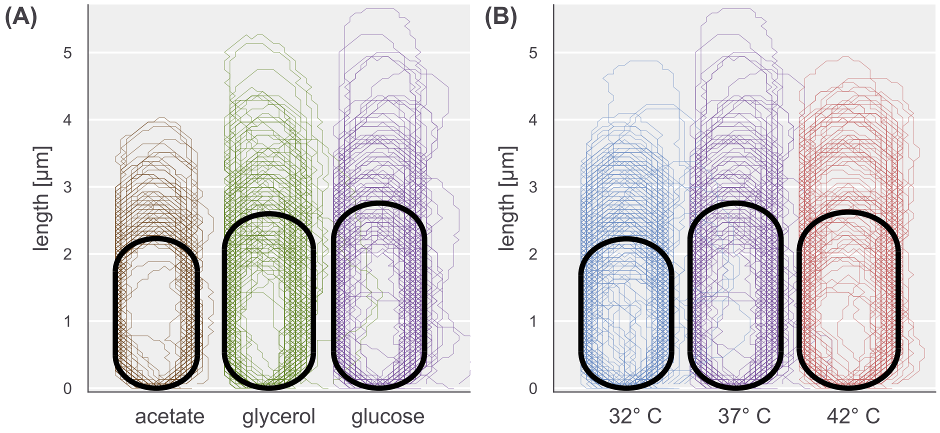

**Fig. S3. Growth-rate dependence of cell shape and spherocylinder approximation.** Contours of segmentation masks for a single experiment of each condition are shown as thin colored lines. Solid lines indicate the approximation of the shape as a spherocylinder given the average length and width of the segmentation masks. Results shown are for (A) carbon quality variation and (B) temperature variation.

### 4. Counting Repressors

In this section, we expand upon the theoretical and experimental implementation of the fluorescence calibration method derived in Ref. (15). We cover several experimental data validation steps as well as details regarding the parameter inference. Finally, we comment on the presence of a systematic error in the repressor counts due to continued asynchronous division between sample preparation and imaging.

**A. Theoretical Background of the Binomial Partitioning Method.** A key component of this work is the direct measurement of the repressor copy number in each growth condition using fluorescence microscopy. To translate between absolute fluorescence and protein copy number, we must be able to estimate the average brightness of a single fluorophore or, in other words, determine a calibration factor $\alpha$ that permits translation from copy number to intensity or vice versa. Several methods have been used over the past decade to estimate this factor, such as measuring single-molecule photobleaching steps (16, 17), measurement of *in vivo* photobleaching rates (18, 19), and through measuring the partitioning of fluorescent molecules between sibling cells after cell division (15, 20, 21). In this work, we used the latter method to estimate the brightness of a single LacI-mCherry dimer. Here, we derive a simple expression which allows the determination of $\alpha$ from measurements of the fluorescence intensities of a collection of sibling cells.

In the absence of measurement error, the fluorescence intensity of a given cell is proportional to the total number of fluorescent proteins $N_{prot}$ by some factor $\alpha$,

$$I_{cell} = \alpha N_{prot}. \quad [13]$$

Assuming that no fluorophores are produced or degraded over the course of the cell cycle, the fluorescent proteins will be partitioned into the two siblings cells such that the intensity of each sibling can be computed as

$$I_1 = \alpha N_1\,;\; I_2 = \alpha(N_{tot} - N_1), \quad [14]$$

where $N_{tot}$ is the total number of proteins in the parent cell and $N_1$ and $N_2$ correspond to the number of proteins in sibling cell 1 and 2, respectively. This explicitly states that fluorescence is conserved upon a division,

$$I_{tot} = I_1 + I_2. \quad [15]$$

As the observed intensity is directly proportional to the number of proteins per cell, measuring the variance in intensity between sibling cells provides some information as to how many proteins were there to begin with. We can compute these fluctuations as the squared intensity difference between the two siblings as

$$\langle (I_1 - I_2)^2 \rangle = \langle (2I_1 - I_{tot})^2 \rangle. \quad [16]$$

We can relate Eq. 16 in terms of the number of proteins using Eq. 13 as

$$\langle (I_1 - I_2)^2 \rangle = 4\alpha^2 \langle N_1^2 \rangle - 4\alpha^2 \langle N_1 \rangle N_{tot} + \alpha^2 N_{tot}^2, \quad [17]$$

where the squared fluctuations are now cast in terms of the first and second moment of the probability distribution for $N_1$.

Without any active partitioning of the proteins into the sibling cells, one can model the probability distribution $g(N_1)$ of finding $N_1$ proteins in sibling cell 1 as a binomial distribution,

$$g(N_1 | \, N_{tot}, p) = \frac{N_{tot}!}{N_1!(N_{tot} - N_1)!} p^{N_1} (1-p)^{N_{tot} - N_1}, \quad [18]$$

where $p$ is the probability of a protein being partitioned into one sibling over the other. With a probability distribution for $N_1$ in hand, we can begin to simplify Eq. 17. Recall that the mean and variance of a binomial distribution are

$$\langle N_1 \rangle = N_{tot} p, \quad [19]$$

and

$$\langle N_1^2 \rangle - \langle N_1 \rangle^2 = N_{tot} p (1-p). \quad [20]$$

With knowledge of the mean and variance, we can solve for the second moment as

$$\langle N_1^2 \rangle = N_{tot} p(1-p) + N_{tot}^2 p^2 = N_{tot} p \left(1 - p + N_{tot} p\right). \quad [21]$$

By plugging Eq. 19 and Eq. 21 into our expression for the fluctuations (Eq. 17), we arrive at

$$\langle (I_1 - I_2)^2 \rangle = 4\alpha^2 \left[ (N_{tot} p[1 - p - N_{tot} p]) - N_{tot}^2 \left( p + \frac{1}{4} \right) \right], \quad [22]$$

which is now defined in terms of the total number of proteins present in the parent cell. Assuming that the proteins are equally partitioned $p = 1/2$, Eq. 22 reduces to

$$\langle (I_1 - I_2)^2 \rangle = \alpha^2 N_{tot} = \alpha I_{tot}. \quad [23]$$

Invoking our assertion that fluorescence is conserved (Eq. 15), $I_{tot}$ is equivalent to the sum total fluorescence of the siblings,

$$\langle (I_1 - I_2)^2 \rangle = \alpha (I_1 + I_2). \quad [24]$$

Thus, given snapshots of cell intensities and information of their lineage, one can compute how may arbitrary fluorescence units correspond to a single fluorescent protein.

**B. Cell Husbandry and Time-Lapse Microscopy.** The fluorescence calibration method derived in Sec. A was first described and implemented by Rosenfeld *et al.* in 2005 (15) followed by a more in-depth approach on the statistical inference of the calibration factor in 2006 (20). In both of these works, the partitioning of a fluorescent protein was tracked across many generations from a single parent cell, permitting inference of a calibration factor from a single lineage. Brewster *et al.* (21) applied this method in a slightly different manner by quantifying the fluorescence across a large number of *single* division events. Thus, rather than examining the partitioning of fluorescence down many branches of a single family tree, it was estimated from an array of single division events where the fluorescence intensity of the parent cell was variable. In the present work, we take a similar approach to that of Brewster *et al.* and examine the partitioning of fluorescence among a large number of independent cell divisions.

A typical experimental work-flow is shown in Fig. S4. For each experiment, the strains were grown in varying concentrations of ATC to tune the expression of the repressor. Once the cells had reached exponential phase growth ($OD_{600nm} \approx 0.3$), the cells were harvested and prepared for imaging. This involved two separate sample handling procedures, one for preparing samples for lineage tracking and estimation of the calibration factor and another for taking snapshots of cells from each ATC induction condition for the calculation of fold-change.

To prepare cells for the calibration factor measurement, a 100 $\mu$L aliquot of each ATC induction condition was combined and mixed in a 1.5 mL centrifuge tube. This cell mixture was then pelleted at 13000$\times g$ for 1 – 2 minutes. The supernatant containing ATC was then aspirated and the pellet was resuspended in an equal volume of sterile growth media without ATC. This washing procedure was repeated three times to ensure that any residual ATC had been removed from the culture and that expression of LacI-mCherry had ceased. Once washed and resuspended, the cells were diluted ten fold into sterile M9 medium and then imaged on a rigid agarose substrate. Depending on the precise growth condition, a variety of positions were imaged for 1.5 to 4 hours with a phase contrast image acquired every 5 to 15 minutes to facilitate lineage tracking. On the final image of the experiment, an mCherry fluorescence image was acquired of every position. The experiments were then transferred to a computing cluster and the images were computationally analyzed, as described in the next section.

During the washing steps, the remaining ATC induced samples were prepared for snapshot imaging to determine the repressor copy number and fold-change in gene expression for each ATC induction condition. Without mixing the induction conditions together, each ATC induced sample was diluted 1:10 into sterile M9 minimal medium and vigorously mixed. Once mixed, a small aliquot of the samples were deposited onto rigid agarose substrates for later imaging. While this step of the experiment was relatively simple, the total preparation procedure typically lasted between 30 and 60 minutes. As is discussed later, the continued growth of the asynchronously growing culture upon dilution into the sterile medium results in a systematic error in the calculation of the repressor copy number.

**C. Lineage Tracking and Fluorescence Quantification.** Segmentation and lineage tracking of both the fluorescence snap shots and time-lapse growth images were performed using the SuperSegger `v1.03` (14) software using MATLAB R2017B (MathWorks, Inc). The result of this segmentation is a list of matrices for each unique imaged position with identifying data for each segmented cell such as an assigned ID number, the ID of the sibling cell, the ID of the parent cell, and various statistics. These files were then analyzed using Python 3.7. All scripts and software used to perform this analysis can be found on the paper website and GitHub repository.

Using the ID numbers assigned to each cell in a given position, we matched all sibling pairs present in the last frame of the growth movie when the final mCherry fluorescence image was acquired. These cells were then filtered to exclude segmentation artifacts (such as exceptionally large or small cells) as well as any cells which the SuperSegger software identified as having an error in segmentation. Given the large number of cells tracked in a given experiment, we could not manually correct these segmentation artifacts, even though it is possible using the software. To err on the side of caution, we did not consider these edge cases in our analysis.

With sibling cells identified, we performed a series of validation checks on the data to ensure that both the experiment and analysis behaved as expected. Three validation checks are illustrated in Fig. S5. To make sure that the computational pairing of sibling cells was correct, we examined the intensity distributions of each sibling pair. If siblings were being paired solely on their lineage history and not by other features (such as size, fluorescence, etc), one would expect the fluorescence distributions between the two sibling cells to be identical. Fig. S5(A) shows the nearly identical intensity distributions of all siblings from a single experiment, indicating that the pairing of siblings is independent of their fluorescence.

Furthermore, we examined the partitioning of the fluorescence between the siblings. In Eq. 19, we defined the mean number of proteins inherited by one sibling. This can be easily translated into the language of intensity as

$$\langle I_1 \rangle = \alpha \left( I_1 + I_2 \right) p, \tag{25}$$

where we assume that fluorescence is conserved during a division event such that $I_{tot} = I_1 + I_2$. In Sec. A, we make the simplifying assumption that partitioning of the fluorophores between the two sibling cells is fair, meaning that $p = 0.5$. We can see if this approximation is valid by computing the fractional intensity of each sibling cell as

$$p = \frac{\langle I_1 \rangle}{\langle I_1 + I_2 \rangle}. \tag{26}$$

Fig. S5(B) shows the distribution of the fractional intensity for each sibling pair. The distribution is approximately symmetric about 0.5, indicating that siblings are correctly paired and that the partitioning of fluorescence is approximately equal between siblings. Furthermore, we see no correlation between the cell volume immediately after division and the observed fractional intensity. This suggests that the probability of partitioning to one sibling or the other is not dependent on the cell size. An assumption [backed by experimental measurements (1, 22)] in our thermodynamic model is that all repressors in a given cell are bound to the chromosome, either specifically or nonspecifically. As the chromosome is duplicated and partitioned into the two siblings without fail, our assumption of repressor adsorption implies that partitioning should be independent of the size of the respective sibling. The collection of these validation statistics give us confidence that both the experimentation and the analysis are properly implemented and not introducing bias into our estimation of the calibration factor.

**D. Statistical Inference of the Fluorescence Calibration Factor.** As is outlined in the Materials and Methods section of the main text, we took a Bayesian approach towards our inference of the calibration factor given fluorescence measurements of sibling cells. Here, we expand in detail on this statistical model and its implementation.

To estimate the calibration factor $\alpha$ from a set of lineage measurements, we assume that fluctuations in intensity resulting from measurement noise is negligible compared to that resulting from binomial partitioning of the repressors upon cell division. In the absence of measurement noise, the intensity of a given cell $I_{cell}$ can be directly related to the total number of fluorophores $N$ through a scaling factor $\alpha$, such that

$$I_{cell} = \alpha N. \qquad [27]$$

Assuming no fluorophores are produced or degraded over the course of a division cycle, the fluorescence of the parent cell before division is equal to the sum of the intensities of the sibling cells,

$$I_{parent} = I_1 + I_2 = \alpha\,(N_1 + N_2)\,, \qquad [28]$$

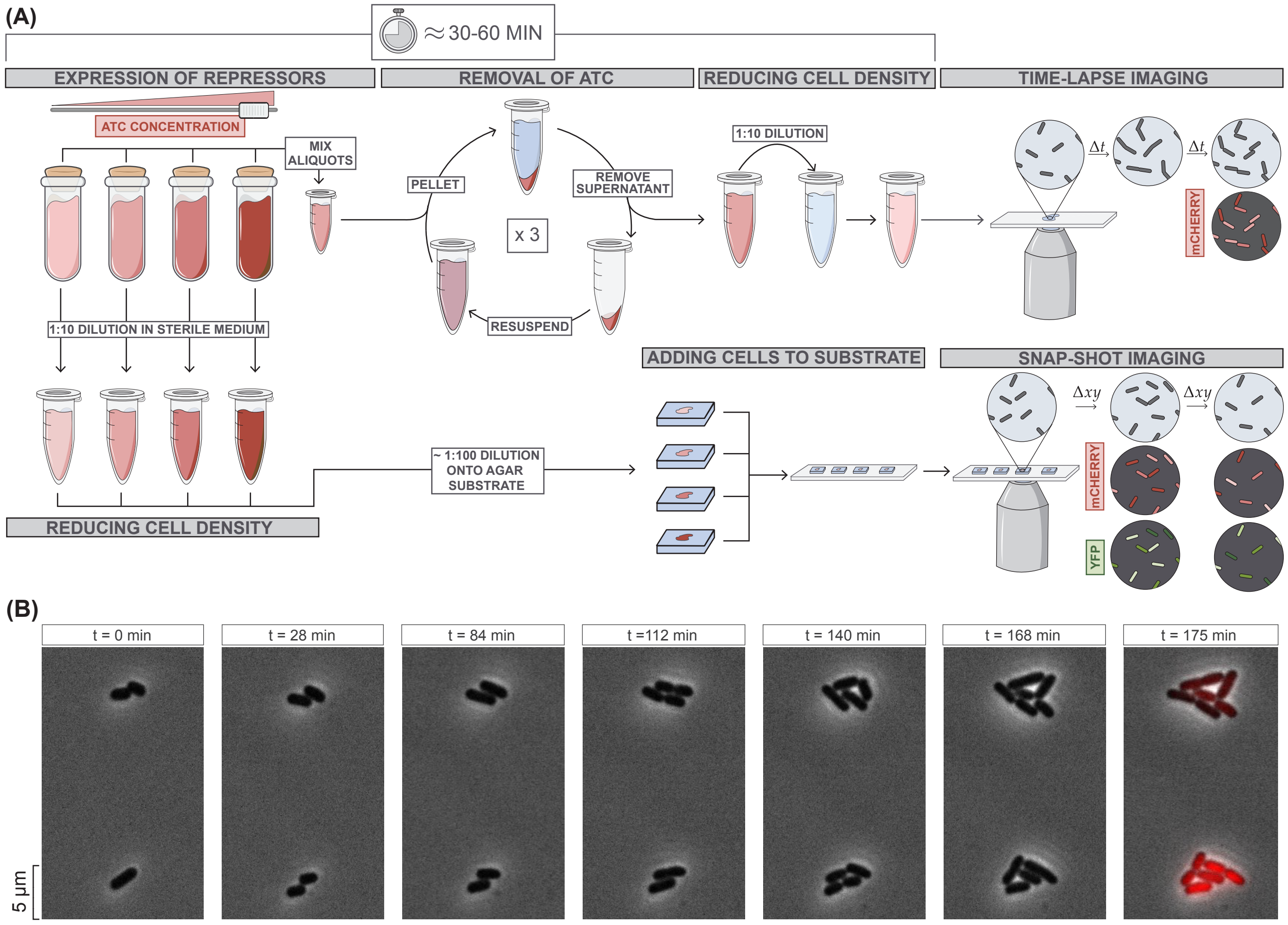

**Fig. S4. Experimental workflow for time-lapse imaging.** (A) The series of steps followed in a given experiment. Cells are grown in varying concentrations of ATC (shaded red cultures) to an $OD_{600nm} \approx 0.3$. Equal aliquots of each ATC-induced culture are mixed into a single eppendorf tube and pelleted via centrifugation. The supernatant containing ATC is aspirated and replaced with an equal volume of sterile, ATC-free growth medium. This washing procedure is repeated three times to ensure that residual ATC is removed from the culture and expression of the LacI-mCherry fusion is ceased. After a final resuspension in sterile ATC-free medium, the cell mixture is diluted 10 fold to reduce cell density. A small aliquot of this mixture is then mounted and imaged at 100x magnification until at least one cell division has occurred. (B) Two representative microcolonies from a time-lapse growth experiment. The time point is provided above each image. After at least one division has occurred, a final mCherry flurorescence image is acquired and quantified.

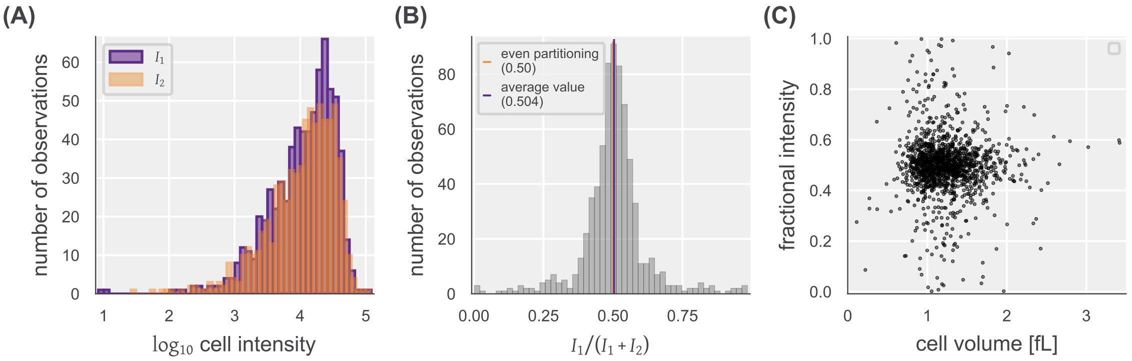

**Fig. S5. Experimental sanity checks and inference of a fluorescence calibration factor.** (A) Intensity distributions of sibling cells after division. Arbitrarily labeled "sibling 1" and "sibling 2" distributions are shown in purple and orange, respectively. Similarity of the distributions illustrates lack of intensity bias on sibling pair assignment. (B) The fractional intensity of sibling 1 upon division. For each sibling pair, the fractional intensity is computed as the intensity of sibling 1 $I_1$ divided by the summed intensities of both siblings $I_1 + I_2$. (C) Partitioning intensity as a function of cell volume. The fractional intensity of every sibling cell were plotted against its estimated newborn volume.

where subscripts 1 and 2 correspond to arbitrary labels of the two sibling cells. We are ultimately interested in knowing the probability of a given value of $\alpha$ which, using Bayes' theorem, can be written as

$$g(\alpha \mid I_1, I_2) = \frac{f(I_1, I_2 \mid \alpha) g(\alpha)}{f(I_1, I_2)}, \tag{29}$$

where we have used $g$ and $f$ to denote probability densities over parameters and data, respectively. The first quantity in the numerator $f(I_1, I_2 \mid \alpha)$ describes the likelihood of observing the data $I_1, I_2$ given a value for the calibration factor $\alpha$. The term $g(\alpha)$ captures all prior knowledge we have about what the calibration factor could be, remaining ignorant of the collected data. The denominator $f(I_1, I_2)$ is the likelihood of observing our data $I_1, I_2$ irrespective of the calibration factor and is a loose measure of how well our statistical model describes the data. As it is difficult to assign a functional form to this term and serves as a multiplicative constant, it can be neglected for the purposes of this work.

Knowing that the two observed sibling cell intensities are related, conditional probability allows us to rewrite the likelihood as

$$f(I_1 \, I_2 \mid \alpha) = f(I_1 \mid I_2, \alpha) f(I_2 \mid \alpha), \tag{30}$$

where $f(I_2 \mid \alpha)$ describes the likelihood of observing $I_2$ given a value of $\alpha$. As $I_2$ can take any value with equal probability, this term can be treated as a constant. Through change of variables and noting that $I_2 = \alpha N_2$, we can cast the likelihood $f(I_1 \mid I_2, \alpha)$ in terms of the number of proteins $N_1$ and $N_2$ as

$$f(I_1 \mid I_2, \alpha) = f(N_1 \mid N_2, \alpha) \left| \frac{dN_1}{dI_1} \right| = \frac{1}{\alpha} f(N_1 \mid N_2, \alpha). \tag{31}$$

Given that the proteins are binomially distributed between the sibling cells with a probability $p$ and that the intensity is proportional to the number of fluorophores, this likelihood becomes

$$f(I_1 \mid I_2, \alpha, p) = \frac{1}{\alpha} \frac{\left(\frac{I_1 + I_2}{\alpha}\right)!}{\left(\frac{I_1}{\alpha}\right)! \left(\frac{I_2}{\alpha}\right)!} p^{\frac{I_1}{\alpha}} (1-p)^{\frac{I_2}{\alpha}} \tag{32}$$

However, The quantity $I/\alpha$ is not exact, making calculation of its factorial undefined. These factorials can therefore be approximated by a gamma function as $n! = \Gamma(n+1)$.

Assuming that partitioning of a protein between the two sibling cells is a fair process ($p = 1/2$), Eq. 32 can be generalized to a set of lineage measurements $[I_1, I_2]$ as

$$f([I_1] \mid [I_2], \alpha) = \frac{1}{\alpha^k} \prod_i^k \frac{\Gamma\left(\frac{I_1 + I_2}{\alpha} + 1\right)}{\Gamma\left(\frac{I_1}{\alpha} + 1\right) \Gamma\left(\frac{I_2}{\alpha} + 1\right)} 2^{-\frac{I_1 + I_2}{\alpha}}, \tag{33}$$

where $k$ is the number of division events observed.

With a likelihood in place, we can now assign a functional form to the prior distribution for the calibration factor $g(\alpha)$. Though ignorant of data, the experimental design is such that imaging of a typical highly-expressing cell will occupy 2/3 of the dynamic range of the camera. We can assume it's more likely that the calibration factor will be closer to 0 a.u. than the bit depth of the camera (4095 a.u.) or larger. We also know that it is physically impossible for the fluorophore to be less than 0 a.u., providing a hard lower-bound on its value. We can therefore impose a weakly informative prior distribution as a half normal distribution,

$$g(\alpha) = \sqrt{\frac{2}{\pi \sigma^2}} \exp\left[\frac{-\alpha^2}{2\sigma^2}\right] \; ; \forall \alpha > 0. \tag{34}$$

where the standard deviation is large, for example, $\sigma = 500$ a.u. / fluorophore. We evaluated the posterior distribution using Markov chain Monte Carlo (MCMC) as is implemented in the Stan probabilistic programming language (23). The `.stan` file associated with this model along with the Python code used to execute it can be accessed on the paper website and GitHub repository. Fig. S6 shows the posterior probability distributions of the calibration factor estimated for several biological replicates of the glucose growth condition at 37° C. For each posterior distribution, the mean and standard deviation was used as the calibration factor and uncertainty for the corresponding data set.

**E. Correcting for Systematic Experimental Error.** While determination of the calibration factor relies on time-resolved measurement of fluorescence partitioning, we computed the repressor copy number and fold-change in gene expression from still snapshots of each ATC induction condition and two control samples, as is illustrated in Fig. S4. Given these snapshots, individual cells were segmented again using the SuperSegger software in MATLAB R2017b. The result of this analysis is an array of single-cell measurements of the YFP and mCherry fluorescence intensities. With these values and a calibration factor estimated by Eq 33 and Eq. 34, we can compute the estimated number of repressors per cell in every condition.

However, as outlined in Sec. B and in Fig. S4, the sample preparation steps for these experiments involve several steps which require careful manual labor. This results in an approximately 30 to 60 minute delay from when production of the LacI-mCherry construct is halted by removal of ATC to actual imaging on the microscope. During this time, the diluted cultures

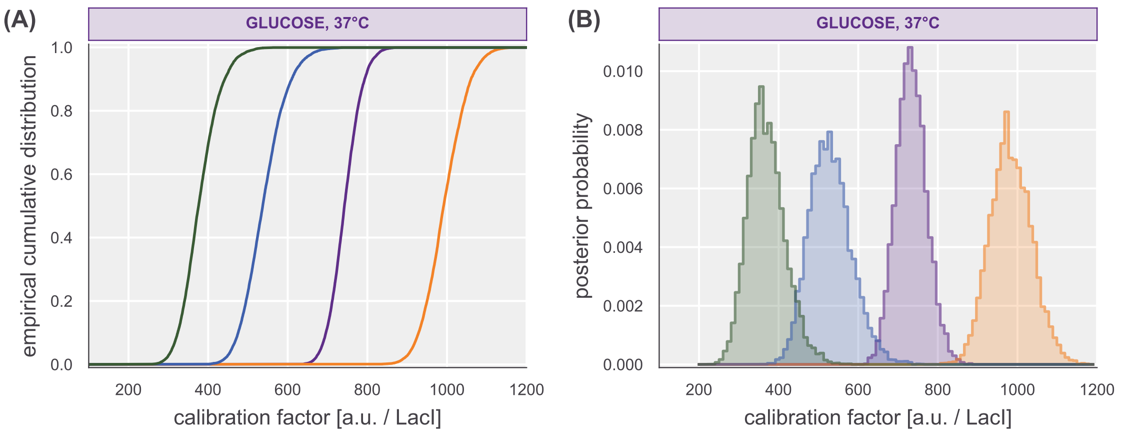

**Fig. S6. Representative posterior distributions for the calibration factor.** (A) Empirical cumulative distribution functions and (B) histograms of the posterior distribution for the calibration factor $\alpha$ for several biological replicates. Different colors indicate different biological replicates of experiments performed in glucose supplemented medium at 37° C.

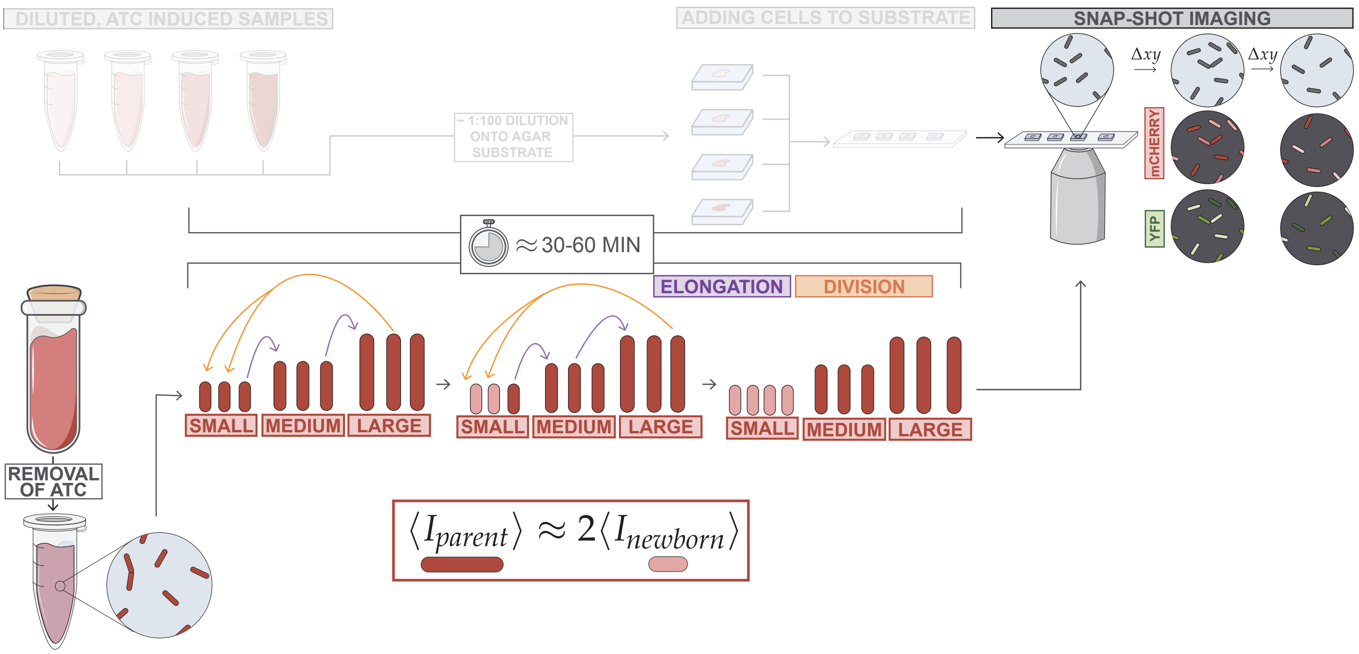

**Fig. S7. Continued division results in a systematic error in repressor counts.** During the sample preparation steps, the asynchronous culture continues to divide even though the length of the sample preparation step is less than the cell doubling time. Because production of LacI-mChery is halted, cells that complete a division cycle during this time will partition their repressors between the progeny. The total number of repressors in parent cells just before division is, on average, twice that of the newborn cells.

are asynchronously growing, meaning that the cells of the culture are at different steps in the cell cycle. Thus, at any point in time, a subset of cells will be on the precipice of undergoing division, partitioning the cytosolic milieu between the two progeny. As LacI-mCherry is no longer being produced, the cells that divided during the dwell time from cell harvesting to imaging will have reduced the number of repressors by a factor of 2 on average. This principle of continued cell division is shown Fig. S7.

How does this partitioning affect our calculation and interpretation of the fold-change in gene expression? Like the LacI-mCherry fusion, the YFP reporter proteins are also partitioned between the progeny after a division event such that, on average, the total YFP signal of the newborn cells is one-half that of the parent cell. As the maturation time of the mCherry and YFP variants used in this work are relatively long in *E. coli* (24, 25), we can make the assumption any newly-expressed YFP molecules after cells have divided are not yet visible in our experiments. Thus, the fold-change in gene expression of the average parent cell can be calculated given knowledge of the average expression of the progeny.

The fold-change in gene expression is a relative measurement to a control which is constitutively expressing YFP. As the latter control sample is also asynchronously dividing, the measured YFP intensity of the newborn constitutively expressing cells is on average 1/2 that of the parent cell. Therefore, the fold-change in gene expression can be calculated as

$$\langle \textit{fold-change}_{parent} \rangle = \frac{2 \times \langle I^{(YFP)}_{newborn}(R > 0) \rangle}{2 \times \langle I^{(YFP)}_{newborn}(R = 0) \rangle} = \frac{\langle I^{(YFP)}_{newborn}(R > 0) \rangle}{\langle I^{(YFP)}_{newborn}(R = 0) \rangle}. \quad [35]$$

Thus, when calculating the fold-change in gene expression one does not need to correct for any cell division that occurs between sample harvesting and imaging as it is a relative measurement. However, the determination of the repressor copy number is a *direct* measurement and requires a consideration of unknown division events.

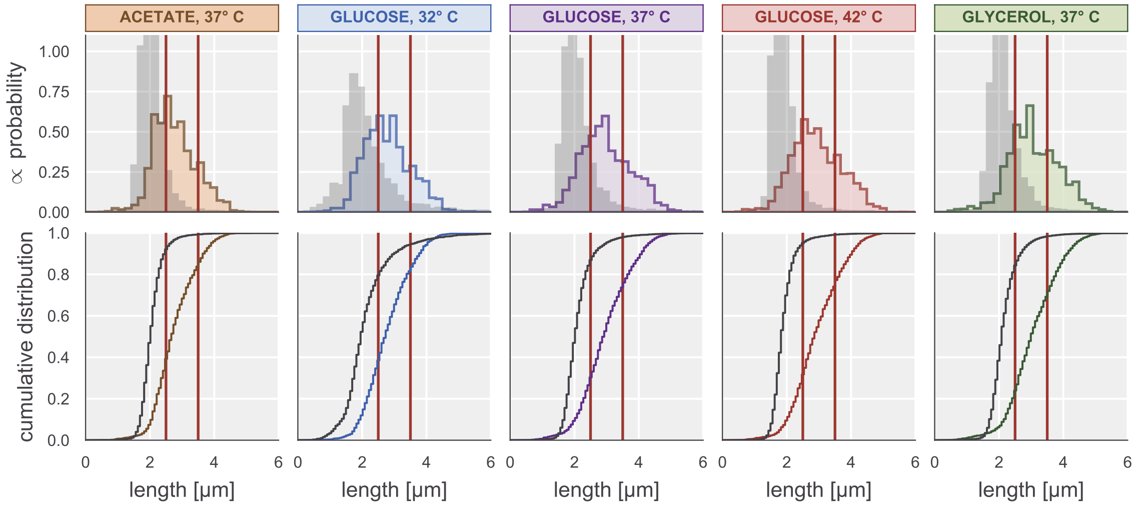

**Fig. S8. Cell length distributions of fluorescence snapshots and newborn cells.** Top row shows the distribution of cell lengths (pole-to-pole, colored distributions) of cells imaged for calculation of the repressor copy number and fold-change in gene expression. The distribution of newborn cell lengths for that given condition is shown in grey. The vertical red lines correspond to the the cell length threshold of 2.5 $\mu$m and 3.5 $\mu$m, from left to right, respectively. Cells to the left of the first vertical line were identified as "small", cells in between the two vertical lines to be "medium" sized, and "long" cells to the right of the second vertical line. Cells below the 2.5 $\mu$m threshold were treated as cells who divided after production of LacI-mCherry had been halted. Bottom row shows the same data as the top row but as the empirical cumulative distribution.

To address this source of systematic experimental error, we examined the cell length distributions of all segmented cells from the snapshots as well as the distribution of newborn cell lengths from the time-lapse measurements. Fig. S8 shows that a significant portion of the cells from the snapshots (colored distributions) overlap with the distribution of newborn cell lengths (grey distributions). For each condition, we partitioned the cells from the snapshots into three bins based on their lengths – "small" cells had a cell length less than 2.5 $\mu$m, "medium" cells had lengths between 2.5 $\mu$m and 3.5 $\mu$m, and "large" cells being longer than 3.5 $\mu$m. These thresholds were chosen manually by examining the newborn cell-size distributions and are shown as red vertical lines in Fig. S8. Under this partitioning, we consider all "small" cells to have divided between cessation of LacI-mCherry production and imaging, "medium"-length cells to have a mixture of long newborn cells (from the tail of the newborn cell length distribution) and cells that haven't divided, and cells in the "long" group to be composed entirely of cells which did not undergo a division over the course of sample preparation.

Given this coarse delineation of cell age by length, we examined how correction factors could be applied to correct for the the undesired systematic error due to dilution of repressors. We took the data collected in this work and compared the results to the fold-change in gene expression reported in the literature for the same regulatory architecture. Without correcting for undesired cell division, the observed fold-change in gene expression falls below the prediction and does not overlap with data from the literature [Fig. S9(A), light purple]. Using the uncorrected measurements, we estimated the DNA binding energy to be $\Delta\varepsilon_R \approx -15\, k_BT$ which does not agree with the value for the O2 operator reported in Garcia and Phillips 2011 (1) or with the inferred DNA binding energies from the other data sources [Fig. S9 (B)]. We direct the reader to Sec. 6 for a more detailed discussion on this parameter inference.

These results emphasize the need to correct for undesired dilution of repressors through cell division during the sample preparation period, and we now consider several different manners of applying this correction. We first consider the extreme case where all cells of the culture underwent an undesired division after LacI-mCherry production was halted. This means that the average repressor copy number measured from all cells is off by a factor of 2. The result of applying a factor of 2 correction to all measurements can be seen in Fig. S9 as dark red points. Upon applying this correction, we find that the observed fold-change in gene expression agrees with the prediction and data from other sources in the literature. The estimated DNA binding energy $\Delta\varepsilon_R$ from these data is also in agreement with other data sources [Fig. S9(B), dark red]. This result suggests that over the course of sample preparation, a non-negligible fraction of the diluted culture undergoes a division event before being imaged.

We now begin to relax assumptions as to what fraction of the measured cells underwent a division event before imaging. As described above, in drawing distinctions between "small", "medium", and "large" cells, we assume the latter represent cells which *did not* undergo a division between the harvesting and imaging of the samples. Thus, the repressor counts of these cells should require no correction. The white-faced points in Fig. S9(A) shows the fold-change in gene expression of *only* the large cell fraction, which falls within error of the theoretical prediction. Furthermore, the inferred DNA binding energy falls within error of that inferred from data of Garcia and Phillips 2011 (1) and that inferred from data assuming all cells underwent a division event [Fig. S9(B)], though it does not fall within error of the binding energy reported in Ref. (1).

The most realistic approach that can be taken to avoid using only the "large" bin of cells is to to assume that all cells with a length $\ell < 2.5\,\mu$m have undergone a division, requiring a two-fold correction to their average repressor copy number. The result of this approach can be seen in Fig. S9 as dark purple points. The inferred DNA binding energy from this correction approach falls within error of that inferred from only the large cells [white-faced points in Fig. S9(B)] as well as overlapping with the estimate treating all cells as having undergone a division (dark red).

There are several experimental techniques that could be implemented to avoid needing to apply a correction factor as described here. In Brewster *et al.* 2014 (21), the fold-change in gene expression was measured by tracking the production rate of a YFP reporter before and after a single cell division event coupled with direct measurement of the repressor copy number using the same binomial partitioning method. This implementation required an extensive degree of manual curation of segmentation as well as correcting for photobleaching of the reporter, which in itself is a non-trivial correction (16). The experimental approach presented here sacrifices a direct measure of the repressor copy number for each cell via the binomial partitioning method, but permits the much higher throughput needed to assay the variety of environmental conditions. Ultimately, the inferred DNA binding energy for all of the scenarios described above agree within $1\,k_BT$, a value smaller than the natural variation in the DNA binding energies of the three native *lacO*, which is $\approx 6\,k_BT$. For the purposes of this work, we erred on the side of caution and only used the cells deemed "large" for the measurements reported in the main text and in the following sections of the SI text.

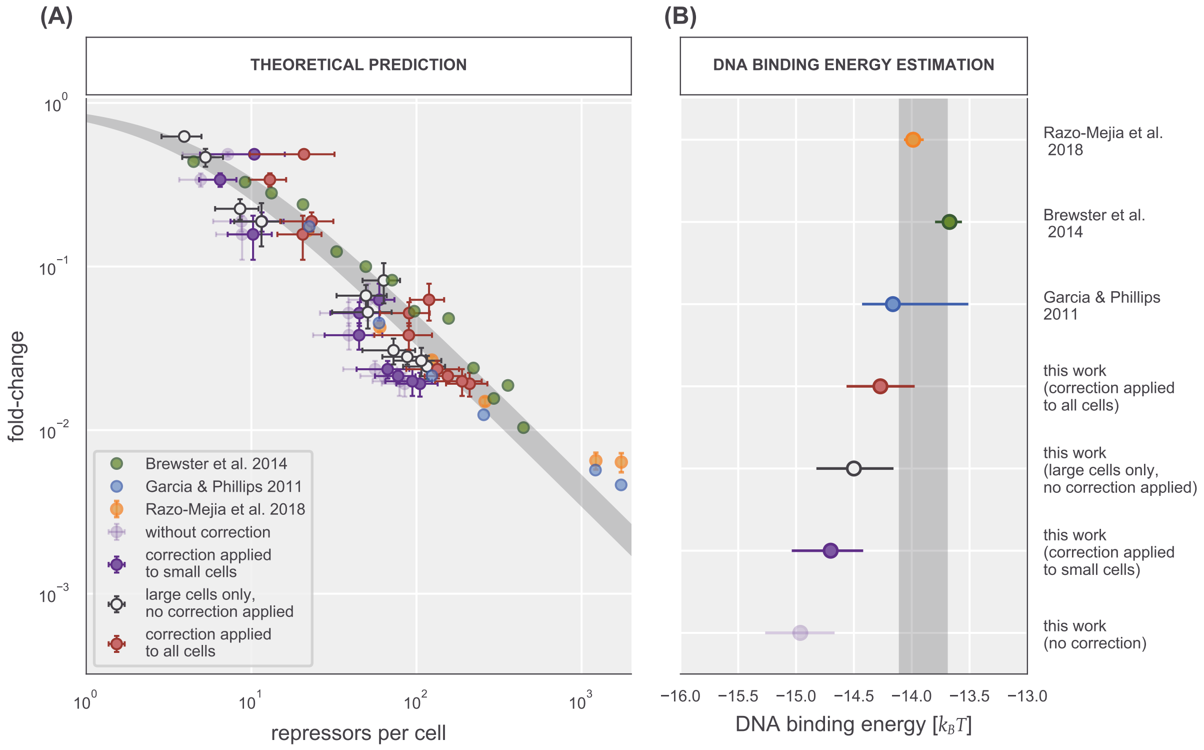

**Fig. S9. Influence of correction factor on fold-change and the DNA binding energy** (A) Fold-change in gene expression measurements from Refs. (1, 2, 21, 26) along with data from this work. Data from this work shown are with no correction (light purple), correcting for small cells only (dark purple), using only the large cell fraction (white-faced points), and treating all cells as newborn cells (dark red). Where visible, errors correspond to the standard error of 5 to 10 biological replicates. (B) Estimated DNA binding energy from each data set. Points are the median of the posterior distribution over $\Delta\varepsilon_R$. Horizontal lines indicate the width of the 95% credible region of the posterior distribution. Grey lines in (A) and (B) correspond to the theoretical prediction and estimated binding energy from Garcia and Phillips 2011, respectively.

### 5. Empirical Determination of the Free Energy Shift $\Delta F$

In the main text, we used the shift in free energy $\Delta F$ as a means to identify how robust the values of the biophysical parameters are to changing conditions, either by changing the carbon source or the temperature of the growth medium. Here, we elaborate on how the observed change in free energy is inferred from measurements of the fold-change in gene expression. We direct the reader to Chure *et al.* 2019 (26) for a more detailed discussion of this inference.

As is given in Eq. 3 of the main text, the fold-change in gene expression can be rewritten in the form of a Fermi function as

$$\textit{fold-change} = \frac{1}{1 + e^{-\beta F}}, \tag{36}$$

where $F$ is the effective free energy difference between the repressor bound and repressor unbound states of the promoter. $F$ can be calculated analytically and has the form

$$F = k_B T \left[\log p_{act}(c) - \log \frac{R}{N_{NS}} + \frac{\Delta\varepsilon_R}{k_B T}\right]. \tag{37}$$

This allows us to draw predictions of the fold-change in gene expression given knowledge of the various biophysical parameter in Eq. 37. However, as dissected in Ref. (26), experimental measurement of the free energy $F$ can be incredibly useful for identifying which parameter may be changing in response to a perturbation.

Given a collection of fold-change measurements, the free energy can be calculated by rearranging Eq. 36 as

$$F = -k_B T \log\left(\frac{1}{\textit{fold-change}} - 1\right). \tag{38}$$

Experimental and statistical noise can push some measurements of the fold-change beyond the theoretical limits with some negative values and others greater than 1. This poses a problem for calculating $F$ directly from the data using Eq. 38 as values below zero or above 1 are undefined.

To confront these boundary conditions, we can model the observed fold-change measurements as being drawn from some distribution whose mean value $\mu$ is constrained to [0, 1], but with a scale parameter $\sigma$ that permits observations beyond. We can then say that the observed free energy is defined by the mean $\mu$ and can be calculated as

$$F = -k_B T \log\left(\frac{1}{\mu} - 1\right) \tag{39}$$

Thus, we wish to estimate the values of $\mu$ and $\sigma$ given a collection of fold-change measurements $\mathbf{fc}$. Using Bayes' theorem, we can calculate this directly as

$$g(\mu, \sigma \mid \mathbf{fc}) = \frac{f(\mathbf{fc} \mid \mu, \sigma) g(\mu, \sigma)}{f(\mathbf{fc})}, \tag{40}$$

where $g$ and $f$ correspond to probability densities over parameters and data respectively. As we did in Sec. A, we can neglect the denominator $f(\mathbf{fc})$ as it serves solely as a normalization factor in this work.

A simple choice for functional form of the likelihood $f(\mathbf{fc} \mid \mu, \sigma)$ [and what was used in Chure et al. 2019 (26)] is a Gaussian distribution,

$$f(\mathbf{fc} \mid \mu, \sigma) = \frac{1}{(2\pi\sigma^2)^{N/2}} \prod_i^N \exp\left[-\frac{(\text{fc}_i - \mu)^2}{2\sigma^2}\right], \tag{41}$$

where $N$ is the total number of measurements in the experiment.

With a likelihood in hand, we can now assign functional forms to the prior distribution $g(\mu, \sigma)$. We begin by making the assumption that as $\sigma$ describes our measurement error, its value should be independent of the distribution mean $\mu$, allowing us to state by conditional probability that

$$g(\mu, \sigma) = g(\mu) g(\sigma). \tag{42}$$

The mean $\mu$ is restricted to the interval [0, 1] as our thermodynamic model forbids fold-change measurements outside of these bounds. We can remain maximally ignorant and say that *any* value between 0 and 1 could be the true value of $\mu$ and assign a prior on this parameter as

$$g(\mu) = \begin{cases} \frac{1}{\mu_{max} - \mu_{min}} & \text{if } \mu_{min} < \mu < \mu_{max} \\ 0 & \text{otherwise} \end{cases}, \tag{43}$$

where the $\mu_{max}$ and $\mu_{min}$ are 1 and 0, respectively.

The standard deviation $\sigma$ permits some measurements to be observed outside of these bounds thereby capturing the experimental noise of the experiments. Given our *a priori* knowledge of the experimental conditions, camera limitations, and other sources of experimental noise, we can say that the value for $\sigma$ is $\leq 1$. As was chosen in Chure *et al.* 2019 (26), we select a half-normal distribution as a prior for $\sigma$,

$$g(\sigma) = \sqrt{\frac{2}{\phi^2 \pi}} \exp\left[-\frac{\sigma^2}{2\phi^2}\right]; \forall \sigma > 0, \tag{44}$$

where $\phi$ is the standard deviation of the half-normal distribution which we choose to be 0.1.

Combining Eq. 41, Eq. 43, and Eq. 44 yields a complete posterior distribution for estimating $\mu$ and $\sigma$ from a collection of fold-change measurements. This posterior was evaluated using Makrov chain Monte Carlo using the Stan probabilistic programming language (23). All Python and Stan code used to perform this inference is available on the paper website and GitHub repository.

### 6. Parameter Estimation of DNA Binding Energies and Comparison Across Carbon Sources

In the main text, we conclude that the biophysical parameters defining the input-output function defined by Eq. 9 are unperturbed between different carbon sources. This conclusion is reached primarily by comparing how well the fold-change and the free energy shift $\Delta F$ is predicted using the parameter values determined in glucose supported medium at 37° C. In this section, we redetermine the DNA binding energy for each carbon source condition and test its ability to predict the fold-change of the other conditions.

**A. Reparameterizing the fold-change input-output function.** As described previously, the fold-change in gene expression is defined by the total repressor copy number $R$, the energetic difference between the active and inactive states of the repressor $\Delta\varepsilon_{AI}$, and the binding energy of the repressor to the DNA $\Delta\varepsilon_R$. Using fluorescence microscopy, we can directly measure the average repressor copy number per cell, reducing the number of variable parameters to only the energetic terms.

Estimating both $\Delta\varepsilon_R$ and $\Delta\varepsilon_{AI}$ simultaneously is fraught with difficulty as the parameters are highly degenerate (2). We can avoid this degeneracy by reparameterizing Eq. 9 as

$$\textit{fold-change} = \left(1 + \frac{R}{N_{NS}} e^{-\beta\epsilon}\right)^{-1}, \quad [45]$$

where $\epsilon$ is the effective energetic parameter

$$\epsilon = \Delta\varepsilon_R - k_B T \log\left(1 + e^{-\beta\Delta\varepsilon_{AI}}\right). \quad [46]$$

Thus, to further elucidate any changes to the parameter values due to changing the carbon source, we can infer the best-fit value of $\epsilon$ for each condition and explore how well it predicts the fold-change in other conditions.

**B. Statistical Inference of $\epsilon$.** We are interested in the probability distribution of the parameter $\epsilon$ given knowledge of the repressor copy number $R$ and a collection of fold-change measurements $\mathbf{fc}$ can be calculated via by Bayes' theorem as

$$g(\epsilon \,|\, R, \mathbf{fc}) = \frac{f(\mathbf{fc} \,|\, R, \epsilon) g(\epsilon)}{f(\mathbf{fc})}, \quad [47]$$

where $g$ and $f$ represent probability densities of parameters and data, respectively. In this context, the denominator term $f(\mathbf{fc})$ serves only as normalization constant and can be neglected. As is described in detail in Refs. (2, 26), we assume that a given set of fold-change measurements are normally distributed about the theoretical value $\mu$ defined by Eq. 9. The likelihood can be mathematically defined as

$$f(\mathbf{fc} \,|\, \epsilon, R) = \frac{1}{(2\pi\sigma^2)^{N/2}} \prod_i^N \exp\left[\frac{-(\text{fc}_i - \mu(\epsilon, R))^2}{2\sigma^2}\right], \quad [48]$$

where $N$ is the total number of fold-change measurements, and $\sigma$ is the standard deviation of the observations about the true mean and is another parameter that must included in the estimation. As the fold-change in gene expression in this work covers several orders of magnitude (from $\approx 10^{-3} - 10^{0}$) it is is better to condition the parameters on the log fold-change rather than linear scaling, translating Eq. 48 to

$$f(\mathbf{fc}^* \,|\, \epsilon, R) = \frac{1}{(2\pi\sigma)^{N/2}} \prod_i^N \exp\left[-\frac{(\text{fc}_i^* - \mu^*(\epsilon, R))^2}{2\sigma^2}\right], \quad [49]$$

where $\mathbf{fc}^*$ and $\mu^*(\epsilon, R)$ are the transformations

$$\mathbf{fc}^* = \log(\mathbf{fc}) \quad [50]$$

and

$$\mu^*(\epsilon, R) = -\log\left(1 + \frac{R}{N_{NS}} e^{-\beta\epsilon}\right), \quad [51]$$

respectively.

With a likelihood in hand, we can now turn towards defining functional forms for the prior distributions $g(\sigma)$ and $g(\epsilon)$. For these definitions, we can turn to those used in Ref. (26),

$$g(\epsilon) \sim \textit{Normal}(\mu = -12, \sigma = 6) \quad [52]$$

and

$$g(\sigma) \sim \textit{HalfNormal}(\phi = 0.1), \quad [53]$$

where we introduce the shorthand notion of "Normal" and "HalfNormal". Combining Eq. 49 and Eqs. 53 – 52 yields the complete posterior distribution for estimating the DNA binding energy for each carbon source medium. The complete posterior distribution was sampled using Markov chain Monte Carlo in the Stan probabilistic programming language (23), whose scripts are available on the paper website and GitHub repository.

The sampled posterior distributions for $\epsilon$ and $\sigma$ for each carbon source condition shown in Fig. S10 and are summarized in Table S1. The posterior distributions of $\epsilon$ across the conditions are approximately equal with highly overlapping 95% credible regions. The predictive capacity of each estimate of $\epsilon$ is shown in Fig. S11 where all fold-change measurements fall upon the theoretical prediction regardless of which carbon source that the parameter value was conditioned upon. With this analysis, we can say with quantitative confidence that the biophysical parameters are indifferent to the physiological changes resulting from variation in carbon quality.

**Table S1. Summarized parameter estimates of $\epsilon$ and $\sigma$ given a single growth condition. Repeated values are the median of the posterior distribution and the upper and lower bounds of the 95% credible region.**

| Growth Condition | Parameter | Value |
|---|---|---|
| Glucose, 37° C | $\epsilon$ | $-14.5^{+0.2}_{-0.3}\, k_BT$ |
| | $\sigma$ | $0.3^{+0.1}_{-0.1}$ |
| Glycerol, 37° C | $\epsilon$ | $-14.6^{+0.1}_{-0.1}\, k_BT$ |
| | $\sigma$ | $0.39^{+0.10}_{-0.08}$ |
| Acetate, 37° C | $\epsilon$ | $-14.1^{+0.2}_{-0.3}\, k_BT$ |
| | $\sigma$ | $0.35^{+0.1}_{-0.1}$ |

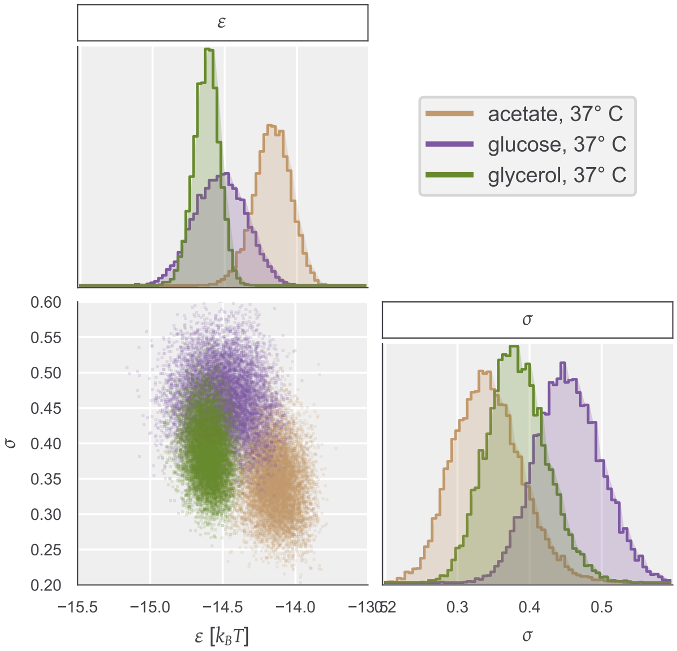

**Fig. S10. Posterior probability distributions of effective DNA binding energy $\epsilon$ and standard deviation $\sigma$.** Top plot shows the marginal probability distribution of $\epsilon$ conditioned only on the glucose (purple), glycerol (green), or acetate (brown) measurements. Bottom left plot shows the joint probability distribution between the effective DNA binding energy $\epsilon$ and the standard deviation $\sigma$. Bottom right shows the marginal posterior distribution over the standard deviation $\sigma$.

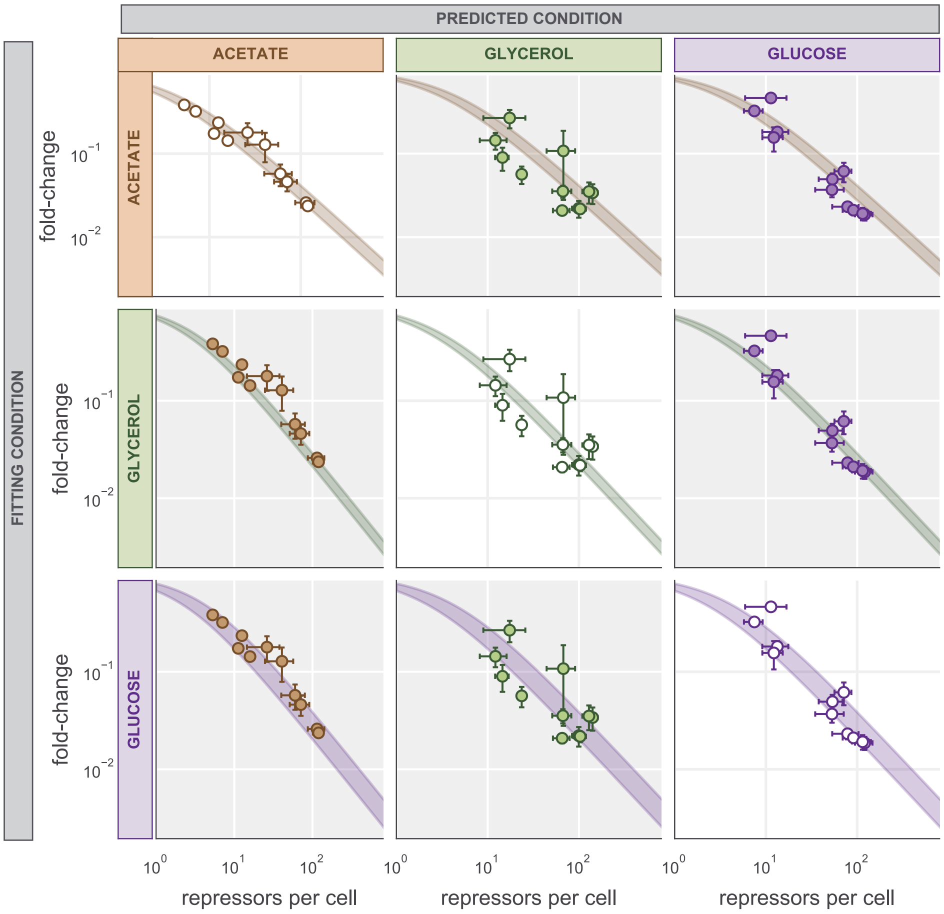

**Fig. S11. Pairwise estimation and prediction of DNA binding energies.** Rows indicate the strain to which the effectice DNA binding energy $\epsilon$ was estimated and columns are the strains whose fold-change is predicted. Shaded lines represent the 95% credible region of the prediction given the estimated value of $\epsilon$. Points and errors correspond to the median and standard error of five to eight biological replicates.

### 7. Statistical Inference of Entropic Costs

In the main text, we describe how a simple rescaling of the energetic parameters $\Delta\varepsilon_R$ and $\Delta\varepsilon_{AI}$ is not sufficient to describe the fold-change in gene expression when the growth temperature is changed from 37° C. In this section, we describe the inference of hidden entropic parameters to phenomenologically describe the temperature dependence of the fold-change in gene expression.

**A. Definition of hidden entropic costs.** The values of the energetic parameters $\Delta\varepsilon_R$ and $\Delta\varepsilon_{AI}$ were determined in a glucose supplemented medium held at 37° C which we denote as $T_{ref}$. A null model to describe temperature dependence of these parameters is to rescale them to the changed temperature $T_{exp}$ as

$$\Delta\varepsilon^* = \frac{T_{ref}}{T_{exp}}\Delta\varepsilon, \quad [54]$$

where $\Delta\varepsilon^*$ is either $\Delta\varepsilon_R$ or $\Delta\varepsilon_{AI}$. However, we found that this null model was not sufficient to describe the fold-change in gene expression, prompting the formulation of a new phenomenological description.

The thermodynamic model defined by Eq. 9 coarse-grains the regulatory architecture to a two state model, meaning many of the rich features of regulation such as vibrational entropy, the material properties of DNA, and the occupancy of the repressor to the DNA are swept into the effective energetic parameters. As temperature was never perturbed when this model was developed, modeling these features was not necessary. However, we must now return to these features to consider what may be affected.

Without assigning a specific mechanism, we can say that there is a temperature-dependent entropic parameter that was neglected in the estimation of the energetic parameters in Garcia and Phillips 2011 (1) and Razo-Mejia *et al.* 2018 (2). In this case, the inferred energetic parameter $\Delta\varepsilon$ is composed of enthalpic ($\Delta H$) and entropic ($\Delta S$) parameters,

$$\Delta\varepsilon^* = \Delta H - T\Delta S. \quad [55]$$

For a set of fold-change measurements at a temperature $T_{exp}$, we are interested in estimating values for $\Delta H$ and $\Delta S$ for each energetic parameter. Given measurements from Refs. (1, 2), we know at 37°$C$ what $\Delta\varepsilon_{RA}$ and $\Delta\varepsilon_{AI}$ are inferred to be, placing a constraint on the possible values of $\Delta H$ and $\Delta S$,

$$\Delta H_R = \Delta\varepsilon_R + T_{ref}\Delta S_R = T_{ref}\Delta S_R - 13.9\,k_B T, \quad [56]$$

and

$$\Delta H_{AI} = \Delta\varepsilon_{AI} + T_{ref}\Delta S_{AI} = T_{ref}\Delta S_{AI} + 4.5\,k_B T \quad [57]$$

for the DNA binding energy and allosteric state energy difference, respectively.

**Statistical inference of $\Delta S_R$ and $\Delta S_{AI}$.** Given the constraints from Eq. 56 and Eq. 57, we are interested in inferring the entropic parameters $\Delta S_R$ and $\Delta S_{AI}$ given literature values for $\Delta\varepsilon_{RA}$ and $\Delta\varepsilon_{AI}$ and the set of fold-change measurements **fc** at a given temperature $T_{exp}$. The posterior probability distribution for the entropic parameters can be enumerated via Bayes' theorem as

$$g(\Delta S_R, \Delta S_{AI}\,|\,\mathbf{fc}) = \frac{f(\mathbf{fc}\,|\,\Delta S_R, \Delta S_{AI})g(\Delta S_R, \Delta S_{AI})}{f(\mathbf{fc})}, \quad [58]$$

where $g$ and $f$ are used to denote probability densities over parameters and data, respectively. As we have done elsewhere in this SI text, we treat $f(\mathbf{fc})$ as a normalization constant and neglect it in our estimation of $g(\Delta S_R, \Delta S_{AI}\,|\,\mathbf{fc})$. Additionally, as is discussed in detail in Sec. 6, we consider the log fold-change measurements to be normally distributed about a mean $\mathrm{fc}^*$ defined by Eq. 9 and a standard deviation $\sigma$. Thus, the likelihood for the fold-change in gene expression is

$$f(\mathbf{fc}\,|\,\Delta S_R, \Delta S_{AI}, \sigma, T_{exp}) = \frac{1}{(2\pi\sigma^2)^{N/2}}\prod_i^N \exp\left[-\frac{\left[\log \mathrm{fc}_i - \log \mathrm{fc}^*(\Delta S_R, \Delta S_{AI}, T_{exp})\right]^2}{2\sigma^2}\right]. \quad [59]$$

In calculating the mean $\mathrm{fc}^*$, the effective energetic parameters $\Delta\varepsilon_R^*$ and $\Delta\varepsilon_{AI}^*$ can be defined and constrained using Eq.56 and Eq. 57 as

$$\Delta\varepsilon_R^* = \Delta H_R - T_{exp}\Delta S_R = \Delta S_R(T_{ref} - T_{exp}) - 13.9\,k_B T_{ref}, \quad [60]$$

and

$$\Delta\varepsilon_{AI}^* = \Delta H_{AI} - T_{exp}\Delta S_{AI} = \Delta S_{AI}(T_{ref} - T_{exp}) + 4.5\,k_B T_{ref}. \quad [61]$$

As the enthalpic parameters are calculated directly from the constraints of Eq. 56 and 57, we must only estimate three parameters, $\Delta S_R$, $\Delta S_{AI}$, and $\sigma$, each of which need a functional form for the prior distribution.

*A priori*, we know that both $\Delta S_R$ and $\Delta S_{AI}$ must be small because $T_{ref}$ and $T_{exp}$ are defined in K. As these entropic parameters can be either positive or negative, we can define the prior distributions $g(\Delta S_R)$ and $g(\Delta S_{AI})$ as a normal distribution centered at zero with a small standard deviation,

$$g(\Delta S_R) \sim Normal(\mu = 0, \sigma = 0.1), \quad [62]$$

and

$$g(\Delta S_{AI}) \sim Normal(\mu = 0, \sigma = 0.1). \quad [63]$$

As we detailed previously in Sec. A and 6, the standard deviation $\sigma$ can be defined as a half-normal distribution centered at 0 with a small standard deviation $\phi$,

$$g(\sigma) \sim HalfNormal(\phi = 0.1). \quad [64]$$

With the priors and likelihood functions in hand, we sampled the posterior distribution using Markov chain Monte Carlo as implemented in the Stan probabilistic programming language (23). We performed three different estimations – one inferring the parameters using only data at 32° C, one using only data from 42° C, and one using data sets from both temperatures pooled together.

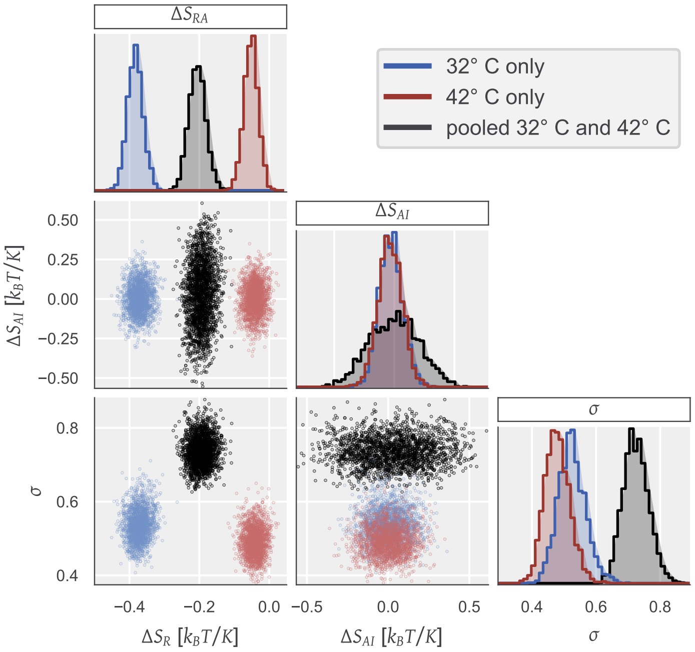

**Fig. S12. Sampled posterior probability distributions of entropic penalty parameter inference**. Marginal and joint distributions conditioned only on data collected at 32° C, only on 42° C, or on both temperatures are shown in blue, red, and black, respectively. The value $\Delta S_R$ and $\Delta S_{AI}$ are given in $k_B T/K$ where $K$ is 1 degree Kelvin.

The sampling results can be seen in Fig. S12. The estimation of $\Delta S_R$ is distinct for each condition where as the sampling for $\Delta S_{AI}$ is the same for all conditions and is centered about 0. The latter suggests that the value of $\Delta\varepsilon_{AI}$ determined at 37° C is not dependent on temperature within the resolution of our experiments. The difference between the estimated value of $\Delta S_R$ between temperatures suggests that there is another component of the temperature dependence that is not captured by the inclusion of a single entropic parameter. Fig. S13 shows that estimating $\Delta S$ from one temperature is not sufficient to predict the fold-change in gene expression at another temperature. The addition of the entropic parameter leads to better fit of the 32° C condition than the simple rescaling of the energy as described by Eq. 54 [Fig. S13, dashed line], but poorly predicts the behavior at 42°C. Performing the inference on the combined 32° C and 42° C data strikes a middle ground between the the predictions resulting from the two temperatures alone [Fig. S13, grey shaded region].

**Entropy as a function of temperature.** In the previous section, we made an approximation of the energetic parameters $\Delta\varepsilon_R$ and $\Delta\varepsilon_{AI}$ to defined by an enthalpic and entropic term, both of which being independent of temperature. However, entropy can be (and in many cases is) dependent on the system temperature, often in a non-trivial manner.

To explore the effects of a temperature-dependent entropy in our prediction of the fold-change, we perform a Taylor expansion of Eq. 55 about the entropic parameter with respect to temperature keeping only the first order term such that

$$\Delta S = \Delta S_0 + \Delta S_1 T, \quad [65]$$

where $\Delta S_0$ is a constant, temperature-independent entropic term and $\Delta S_1$ is the entropic contribution per degree Kelvin. With this simple relationship enumerated, we can now define the temperature-dependent effective free energy parameters $\Delta\varepsilon^*_R$ and $\Delta\varepsilon^*_{AI}$ as

$$\Delta\varepsilon^*_R = (S_{0R} + S_{1R} T_{exp})(T_{ref} - T_{exp}) - 13.9\, k_B T, \quad [66]$$

and

$$\Delta\varepsilon^*_{AI} = (S_{0AI} + S_{1AI} T_{exp})(T_{ref} - T_{exp}) + 4.5\, k_B T, \quad [67]$$

respectively, again relying on the constraints defined by Eq. 60 and Eq. 61.

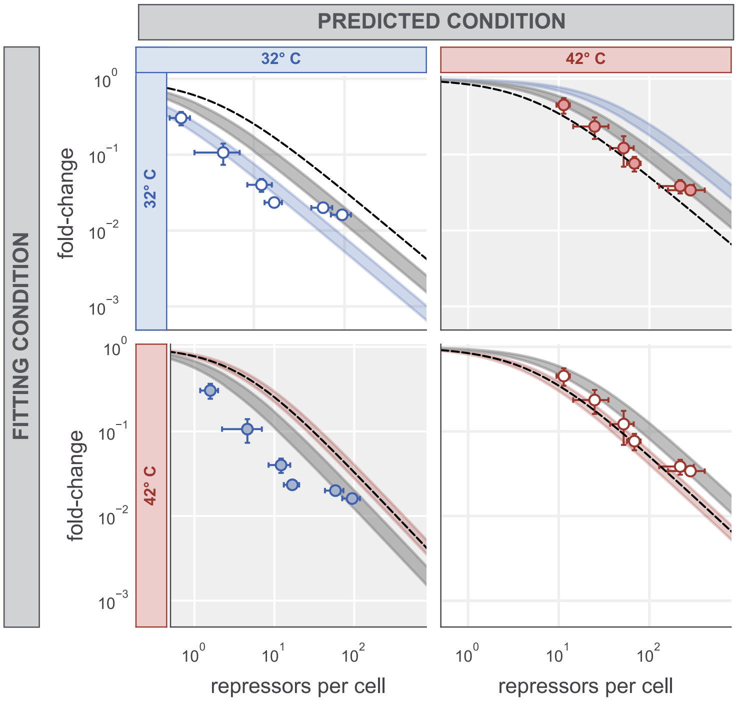

**Fig. S13. Pairwise predictions of fold-change in gene expression at different temperatures.** Each row represents the condition used to infer the entropic parameters and columns are the conditions that are being predicted. Color shaded regions represent the 95% credible region of the predicted fold-change given the inferred values of $\Delta S_R$ and $\Delta S_{AI}$ for each condition. The black dashed line represents the predicted fold-change in gene expression by a simple rescaling of the binding energy determined at 37° C. The grey shaded region is the 95% credible region of the fold-change given estimation of $\Delta S_R$ and $\Delta S_{AI}$ conditioned on both temperatures pooled together.

Using a similar inferential approach as described in the previous section, we sample the posterior distribution of these parameters using Markov chain Monte Carlo and compute the fold-change and shift in free energy for each temperature. As seen in Fig. S14, there is a negligible improvement in the description of the data by including this temperature dependent entropic parameter.

These results together suggest that our understanding of temperature dependence in this regulatory architecture is incomplete and requires further research from both theoretical and experimental standpoints.

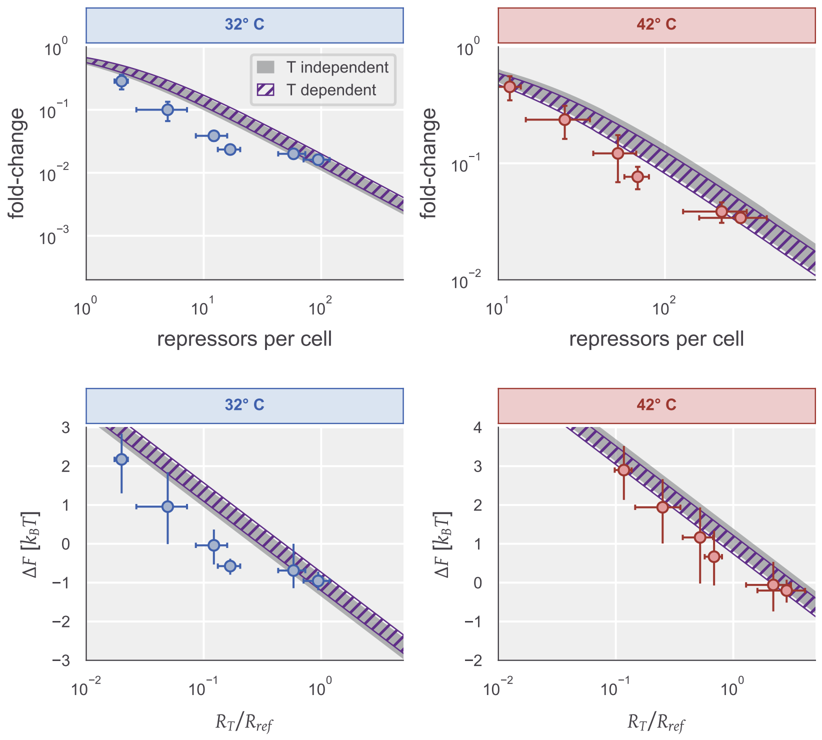

**Fig. S14. Fold-change and shift in free energy including a temperature-dependent entropic contribution.** Top row illustrates the estimated fold-change in gene expression at 32° C (left) and 42° C (right). Bottom row shows the estimated shift in free energy for each temperature. The grey transparent line in all plots shows the relevant quantity including only a temperature-independent entropic parameter. Purple hashed line illustrates the relevant quantity including a temperature-dependent entropic contribution. The width of each curve indicates the 95% credible region of the relevant quantity. Points in each correspond to the experimental measurements. The points in the top row represent the mean and standard error of five to eight biological replicats. Points in the bottom row correspond to the median value of the inferred free energy shift and the mean of 5 to 8 biological replicates for the repressor copy number. Vertical error represent the upper and lower bounds of the 95% credible region of the pararmeter. Horizontal errorbars correspond to the standard error of 5 to 8 biological replicates.

### 8. Media Recipes and Bacterial Strains

The primary interest in varying the available carbon source in growth media was to modulate the quality of the carbon rather than the quantity. With this in mind, we developed the various growth media to contain the same net number of carbon atoms per cell. The standard reference was 0.5% (weight/volume) glucose (1), which results in $10^8$ carbon atoms per $10^{-15}$ L. The base recipe is given in Table S2. The bacterial strains used in this work are given in S3.

**Table S2. M9 minimal medium recipe for each carbon source used in this work.**

| Ingredient | Stock Concentration | Volume | Final Concentration |
|---|---|---|---|
| dd$H_2O$ | – | 773 mL | - |
| $CaCl_2$ | 1 M | 100 $\mu$L | 100 $\mu$M |
| $MgSO_4$ | 1 M | 2 mL | 2 mM |
| M9 Salts (BD Medical, Cat. No. 248510) | 5X | 200 mL | – |
| _Carbon Source_ | | | $10^8$ C / fL |
| Glucose | 20% (weight/volume) | 25 mL | 0.5% (weight/volume) |
| Glycerol | 20% (weight/volume) | 25 mL | 0.5% (weight/volume) |
| Acetate | 20% (weight/volume) | 25 mL | 0.5% (weight/volume) |
| total | | 1 L | 1X |

**Table S3. Bacterial strains used in this work, generated by Ref. (21)**

| Genotype | Plasmid | Notes |
|---|---|---|
| MG1655::$\Delta$lacZYA;_intC<>4*CFP_ | – | Autofluorescence control |
| MG1655::$\Delta$lacZYA;_intC<>4*CFP_;_galK<>25-O2+11-YFP_ | – | Constitutive expression control |
| MG1655::$\Delta$lacZYA;_intC<>4*CFP_;_galK<>25-O2+11-YFP_;_ybcN<>1-lacI($\Delta$353-363)-mCherry_ | pZS3$P_{N25}$-tetR | Strain with ATC inducible LacI-mCherry |

### References

1. Garcia HG, Phillips R (2011) Quantitative dissection of the simple repression input-output function. *Proceedings of the National Academy of Sciences* 108(29):12173–12178.
2. Razo-Mejia M, et al. (2018) Tuning Transcriptional Regulation through Signaling: A Predictive Theory of Allosteric Induction. *Cell Systems* 6(4):456–469.e10.
3. Klumpp S, Hwa T (2014) Bacterial growth: Global effects on gene expression, growth feedback and proteome partition. *Current Opinion in Biotechnology* 28:96–102.
4. Jones DL, Brewster RC, Phillips R (2014) Promoter architecture dictates cell-to-cell variability in gene expression. *Science* 346(6216):1533–1536.
5. Schaechter M, Maaløe O, Kjeldgaard NO (1958) Dependency on Medium and Temperature of Cell Size and Chemical Composition during Balanced Growth of *salmonella Typhimurium*. *Microbiology* 19(3):592–606.
6. Jun S, Si F, Pugatch R, Scott M (2018) Fundamental principles in bacterial physiology—history, recent progress, and the future with focus on cell size control: A review. *Reports on Progress in Physics* 81(5):056601.
7. Allen RJ, Waclaw B (2018) Bacterial growth: A statistical physicist's guide. *Reports on Progress in Physics* 82(1):016601.
8. Swain PS, et al. (2016) Inferring time derivatives including cell growth rates using Gaussian processes. *Nature Communications* 7:13766.
9. Pilizota T, Shaevitz JW (2012) Fast, Multiphase Volume Adaptation to Hyperosmotic Shock by *Escherichia coli*. *PLOS ONE* 7(4):e35205.
10. Pilizota T, Shaevitz JW (2014) Origins of *Escherichia* coli Growth Rate and Cell Shape Changes at High External Osmolality. *Biophysical Journal* 107(8):1962–1969.
11. Taheri-Araghi S, et al. (2015) Cell-size control and homeostasis in bacteria. *Current biology: CB* 25(3):385–391.
12. Schmidt A, et al. (2016) The quantitative and condition-dependent *Escherichia* coli proteome. *Nature Biotechnology* 34(1):104–110.
13. Kubitschek HE, Friske JA (1986) Determination of bacterial cell volume with the Coulter Counter. *Journal of Bacteriology* 168(3):1466–1467.
14. Stylianidou S, Brennan C, Nissen SB, Kuwada NJ, Wiggins PA (2016) SuperSegger: Robust image segmentation, analysis and lineage tracking of bacterial cells. *Molecular Microbiology* 102(4):690–700.
15. Rosenfeld N, Young JW, Alon U, Swain PS, Elowitz MB (2005) Gene Regulation at the Single-Cell Level. *Science* 307(5717):1962–1965.
16. Garcia HG, Lee HJ, Boedicker JQ, Phillips R (2011) Comparison and Calibration of Different Reporters for Quantitative Analysis of Gene Expression. *Biophysical Journal* 101(3):535–544.
17. Bialecka-Fornal M, Lee HJ, DeBerg HA, Gandhi CS, Phillips R (2012) Single-Cell Census of Mechanosensitive Channels in Living Bacteria. *PLoS ONE* 7(3):e33077.
18. Nayak CR, Rutenberg AD (2011) Quantification of Fluorophore Copy Number from Intrinsic Fluctuations during Fluorescence Photobleaching. *Biophysical Journal* 101(9):2284–2293.
19. Kim NH, et al. (2016) Real-time transposable element activity in individual live cells. *Proceedings of the National Academy of Sciences* 113(26):7278–7283.
20. Rosenfeld N, Perkins TJ, Alon U, Elowitz MB, Swain PS (2006) A Fluctuation Method to Quantify In Vivo Fluorescence Data. *Biophysical Journal* 91(2):759–766.
21. Brewster RC, et al. (2014) The Transcription Factor Titration Effect Dictates Level of Gene Expression. *Cell* 156(6):1312–1323.
22. Phillips R, et al. (2019) Figure 1 Theory Meets Figure 2 Experiments in the Study of Gene Expression. *Annual Review of Biophysics* 48(1):121–163.
23. Carpenter B, et al. (2017) Stan: A Probabilistic Programming Language. *Journal of Statistical Software* 76(1):1–32.
24. Balleza E, Kim JM, Cluzel P (2018) Systematic characterization of maturation time of fluorescent proteins in living cells. *Nature methods* 15(1):47–51.
25. Nagai T, et al. (2002) A variant of yellow fluorescent protein with fast and efficient maturation for cell-biological applications. *Nature Biotechnology* 20(1):87–90.
26. Chure G, et al. (2019) Predictive Shifts in Free Energy Couple Mutations to Their Phenotypic Consequences. *Proceedings of the National Academy of Sciences* 116(37).
